# Probabilistic cell lineage inference from mitochondrial mutations

**DOI:** 10.1101/2025.02.28.640740

**Authors:** Sophie Seidel, Joanna Hård, Pedro F. Ferreira, Pietro Berkes, Katharina Jahn, Carl-Johan Eriksson, Jeff E. Mold, Jakob Michaelsson, Jack Kuipers, Niko Beerenwinkel

## Abstract

Tracing genealogical relationships between cells and their molecular states is essential for understanding tissue development, maintenance and disease. We present mt-SCITE, a computational method that reconstructs cell division histories from mitochondrial mutations in single cells. On simulated and experimental benchmarks, mt-SCITE outperformed existing methods in terms of cell lineage reconstruction, and scaled to thousands of cells. Applied to expanded CD8+ T cells following vaccination, mt-SCITE resolved subclonal relationships and revealed clones with predominantly memory-like, effector-like, or mixed states. mt-SCITE robustly identifies clonally related populations and resolves substructure within them, providing a broadly applicable approach to study cell lineage development.

## Introduction

Tissue development, regeneration, and homeostasis are regulated by the growth and development of clonally related cell populations arising from a single ancestor cell. How a particular cell type develops, and what its relationships to progenitor cells are, remain fundamental open questions that may have important implications for how we view the body in health and in disease. Cell fate and clonal dynamics have been studied in model organisms using lineage tracing methods that introduce heritable labels into individual cells such that their clonal progeny can be tracked at later time points. Most lineage tracing approaches rely on engineered genetic tags, including, for example, fluorescent reporter genes, Cre-mediated recombination, or CRISPR-based genetic scars [1, 2]. The recent development of advanced molecular tools has enabled combined approaches for lineage tracing with single-cell RNA sequencing (scRNA-seq) to interrogate both cell lineage relationships and cell states [3, 4]. However, an understanding of cell turnover dynamics, including the origins and regenerative potential of distinct cell types, remains limited in humans, because fate mapping techniques to study cell clonality that involve genetic manipulation can only be applied in experimental animal models [2, 5–10].

The development of single-cell whole-genome DNA sequencing technologies, combined with statistical and computational methods for analyzing somatic mutations, have recently made it possible to infer cell lineage relationships of human cells, leveraging the fact that the inheritance patterns of somatic mutations in cells will mirror their genealogical relationships [10–13]. However, due to the large size of the nuclear genome, single-cell whole-genome sequencing remains an expensive approach and poses technical difficulties, particularly in accurately identifying heterozygous variants present in just one DNA copy per cell [14, 15]. These limitations motivate the exploration of alternative approaches for lineage tracing in humans.

DNA methylation has recently been proposed as an alternative source of lineage information. Stochastic epimutations accumulate at CpG sites with successive cell divisions and, when heritably and neutrally maintained, mark shared ancestry [16]. Two single-cell strategies have been developed. The first, MethylTree [17], relies on genome-wide methylome profiles and to handle the resulting data sparsity computes a cell-cell similarity matrix and iteratively denoises it before identifying clones. The second, EPI-Clone [18], profiles a pre-selected panel of informative CpGs, which lowers the sequencing burden and increases per-site coverage but requires the informative sites to be defined a priori and calibrated for each tissue. This approach recovers clones and has so far been demonstrated in blood. A limitation common to both approaches is that CpG methylation encodes cell-type identity and transcriptional state in addition to ancestry, so lineage-informative sites must be separated from those that track cell state [17, 18]. Mitochondrial mutations are not subject to this confounding, as they act as passenger markers that are largely independent of cell state and can be recovered from the same single-cell assays without a tissue-specific panel.

Beyond avoiding this confounding, mitochondrial mutations offer several further advantages for reconstructing cell lineages. First, mitochondrial mutations may be detected in Assay for Transposase-Accessible Chromatin with high-throughput sequencing (ATAC-seq) and scRNA-seq data [19–26] and are thus already available in existing public datasets. ATAC-seq assesses the potential for genes to be expressed, while scRNA-seq measures gene expression directly. The data generated by these technologies enable accurate classification of cell types [27]. Therefore, if ATAC-seq and scRNA-seq data could be used for lineage tracing, they would have the advantage over DNA sequencing of providing additional information about the function and identity of cells. Second, utilizing mitochondrial variants for cell lineage reconstruction is cost-effective given the small size of the mitochondrial genome of approximately 16.6kb in humans [21, 24, 26]. Third, mutations are abundant as the mutation rate is estimated to be 10 to 100-fold higher in mitochondrial DNA (mtDNA) as compared to the nuclear genome, resulting in faster accumulation of variants and hence stronger phylogenetic signals [21, 28–31]. Finally, mtDNA mutations are detectable due to the high copy number of mtDNA, typically 1,000–10,000 copies per cell, which reduces random drop-out in single-cell amplification [28].

However, mtDNA data differs from other endogenous lineage markers in ways that result in new data analysis challenges. In mtDNA, a de novo variant will arise in only one of the thousands of mtDNA copies. Some variants will go extinct and some will be replicated and rise to a higher frequency in the mtDNA pool. The resulting coexistence of mutant and wildtype mtDNA within a single cell is termed heteroplasmy; we refer to the fraction of copies carrying a given variant as its heteroplasmy frequency (or equivalently its variant allele frequency, VAF). When the heteroplasmy level is still low it may be undetectable or challenging to distinguish from errors introduced during amplification or sequencing. The high mutation rate in mtDNA may also result in mutations appearing independently in unrelated cells (homoplasy). The mitochondrial genome is transcribed in almost its entirety, and therefore, new mitochondrial variants may be subject to selective pressures [21]. Moreover, mitochondrial variation is subject to genetic drift. This may result from mtDNA turnover outside of cell division, a process termed relaxed replication. A second mechanism underlying genetic drift in mtDNA is vegetative segregation, in which mitochondria, and thus mtDNA, may be unevenly distributed into daughter cells during cell division [32]. Taken together, while mitochondrial mutations are associated with several advantages for lineage tracing, their complex dynamics present challenges for cell lineage tree reconstruction.

Computational methods designed for calling mtDNA variants in ATAC-seq and scRNA-seq data have been developed in recent years [19, 20, 23]. Mitochondrial mutations were subsequently used to identify clonal cell populations based on the notion that clonally related cells will share mitochondrial mutations. Clonal cell populations were represented by the presence or absence of detected mitochondrial mutations or identified through hierarchical clustering [20, 21, 23, 26]. The binary representation of presence or absence is limited because it may require stringent filtering to distinguish informative variants, and perceived absence may result from false negative genotypes, for example, due to low read coverage at variant loci. While clustering methods consider the mutation profile to group cells based on pairwise distances, they do not consider any higher order distances, such as the genealogical relationships of cells. Moreover, clustering may perform poorly or fail completely when presented with noisy and inconsistent data as is the case with single-cell data [33].

Recently, MERLIN [34], a method for tree inference based on mitochondrial mutations was introduced. MERLIN employs a matrix decomposition approach to decompose the observed cell-by-mutation frequency matrix into a cell-by-clone matrix and a clone-by-mutation matrix, which can be subsequently interpreted as a clone and mutation tree. However, MERLIN does not model sequencing errors, which are prevalent in current sequencing settings, and makes strong assumptions about mitochondrial dynamics, such as no clone going extinct.

Here, we present an inference framework for cell lineage reconstruction. **M**i**T**ochondrial mutation-based **S**ingle **C**ell **I**nference and **T**ree reconstruction of cell lineage **E**volution (mt-SCITE) models somatic mutation probabilities in mtDNA captured by ATAC-seq and scRNA-seq protocols and reconstructs the cell lineage in form of a mutation tree consisting of acquired mitochondrial mutations with cells attached to them. Provided with a set of putatively informative mutations, mt-SCITE computes per-cell mutation probabilities that account for uncertainty from sequencing errors and low coverage, and uses these to further filter the input set. Subsequently, mt-SCITE searches for the tree that best explains the data across the selected range of sequencing error rates. The method is robust to dropout of mutations because when inferring the mutation tree, it marginalizes each cell’s attachment over all tree positions, so the placement of a mutation reflects the consensus of all cells carrying it rather than any single noisy observation. We compared mt-SCITE against several methods, including Neighbour-joining (NJ), Unweighted pair group method with arithmetic mean (UPGMA), MERLIN, and SCITE, using simulated data and experimental data with a ground truth tree. Further, we evaluated the performance of mt-SCITE in resolving cell lineage relationships on *in vitro* expanded human CD8+ T-cells from ATAC-seq and *in vivo* expanded CD8+ T-cell clones [35–37] from single-cell RNA sequencing data. In the latter dataset, T-cell receptor (TCR) rearrangements were leveraged as natural barcodes for clonality and thus provide the ground truth clonal relationships [21]. Comparing the clones within the tree structure with the ground truth clones determined by the TCR, we found that mt-SCITE robustly enables the identification of human cell clones *in vivo*. Furthermore, our tree inference approach allowed us to detect subclonal structures within the T-cell clones and to unveil the lineage relationships between single cells, increasing the resolution of lineage tracing compared to published methods, which are focused on inferring the lineage relationships between clones [20, 22, 23].

## Results

We introduced mt-SCITE, a computational tool for estimating cell lineage relationships based on mitochondrial mutations, compared its performance in reconstructing the tree with existing methods on simulated data and an experimental dataset with a ground truth lineage tree, and applied it to two datasets of T-cell expansion following vaccination. We further apply mt-SCITE to the MAESTER dataset [22], to demonstrate its scalability to large numbers of cells.

### Phylogenetic inference of cell lineage relationships based on shared mitochondrial mutations

mt-SCITE leverages somatic mitochondrial genetic variation to infer the evolutionary history of dividing cells. In contrast to most existing methods, which take binarised mutation calls as input, mt-SCITE operates directly on raw read counts. This allows it to compute a probabilistic estimate of mutation presence at each position in each cell, and to learn the sequencing error rate that produces the most likely tree on a held-out partition of the data (Methods).

The first step in our workflow is the computation of mutation probabilities, for each sample and for each position in the mitochondrial genome (Figure 1). Throughout, ”sample” denotes the unit attached to the inferred tree. For single-cell assays (scRNA-seq, MAESTER) a sample is one cell; for the bulk ATAC-seq data a sample is a bulk-sequenced clonal population. mt-SCITE treats both identically, since it operates on per-sample read-count matrices regardless of whether those reads come from one cell or many.

**Fig. 1:**
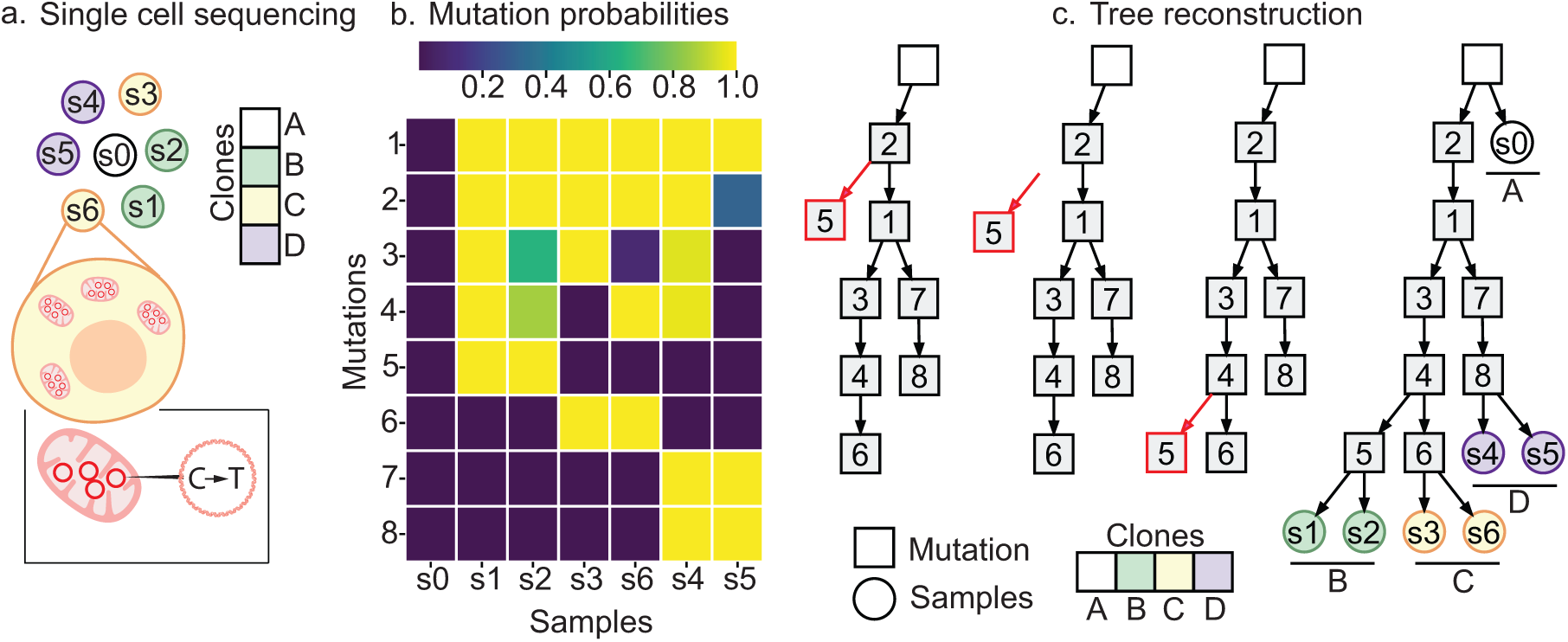
(a) Somatic mitochondrial mutations arising in a cell are passed on to daughter cells during cell division and can be detected by sequencing. (b) From the read counts, mt-SCITE computes the mutation probability matrix, in which each entry is the probability that a given mitochondrial mutations (row) is present in a given sample (column); a sample can be a single cell or a bulk-sequenced clonal population. (c) These mutation probabilities are used to infer the maximum likelihood evolutionary history of the samples. The tree space is explored using a MCMC sampling scheme. During this process, branches are pruned and reattached, as indicated in red, according to the likelihood of the tree. mt-SCITE outputs the trees with the best likelihood. The trees consist of a root node (white square), the acquired mutations (grey squares) and samples (circles) attached to the mutation paths.

The mutation probabilities are computed based on the read counts supporting the four different nucleotides and depend on the error rate of the sequencing experiment (Methods). The higher the fraction of reads for an alternative nucleotide relative to the sequencing error, the higher the probability for a mutation. Given that the error rate of a sequencing experiment is typically unknown in practice, we introduce a procedure for learning the error rate from the data using cross-validation (Methods).

In the second step, the mitochondrial mutation probabilities are provided as input to infer a mutation tree encoding the cell lineage relationships. For this task, we developed a new version of a stochastic search and sampling algorithm for phylogenetic inference extending [38] to handle mitochondrial mutations. We use Markov Chain Monte Carlo (MCMC) to explore the tree space and find the maximum likelihood evolutionary history of the dividing cells. The output of mt-SCITE is a maximum likelihood tree where the nodes represent mutations accumulating over time and the samples are attached to the mutation node best representing their read counts. Samples attached to the root node have no mutations in any of the mutation sites. A sample attached to any other part of the tree has the mutation of the node it is attached to and all the mutations of its ancestors. The tree inference approach shares information between samples, thereby increasing the statistical power to detect mutations as compared to calling mutations independently.

### Benchmark on simulated data

We compare mt-SCITE to the recently published method MERLIN [34], the standard heuristic pipeline of mutation calling followed by distance-based tree reconstruction (NJ [39] and UPGMA [40]), and SCITE [38] as an additional baseline. To compare their performance, we simulated ground truth mutation and cell lineage trees for *L*=10, 50 mutations and *N* =100, 500 cells (Figure 2, Supplemental Figures 5-12). Mitochondrial mutation frequencies were simulated along the mutation trees using an initial mitochondrial variant frequency *f*_init_ = 1%, 10% for newly arising mutations. We then randomly sampled where in the mutation trees cells would attach, which specifies their mutation profile and heteroplasmy frequencies. Based on the heteroplasmy frequencies, we sampled read counts for each cell from a multinomial distribution, incorporating varying sequencing error rates (*ρ* = 0.05, 0.005, 0.0005). SCITE and mt-SCITE were applied with fixed error rates *ρ*. For mt-SCITE, we also evaluated performance with learned error rates. As MERLIN, NJ and UPGMA do not model the sequencing error rates, we report a single performance value for each. Mutation tree accuracy is quantified using the normalised parent-child distance [41] for the methods that produce mutation trees (mt-SCITE, SCITE, MERLIN). Cell lineage accuracy is evaluated for all methods using the root mean squared error (RMSE) between inferred and simulated cell-cell distance matrices.

**Fig. 2:**
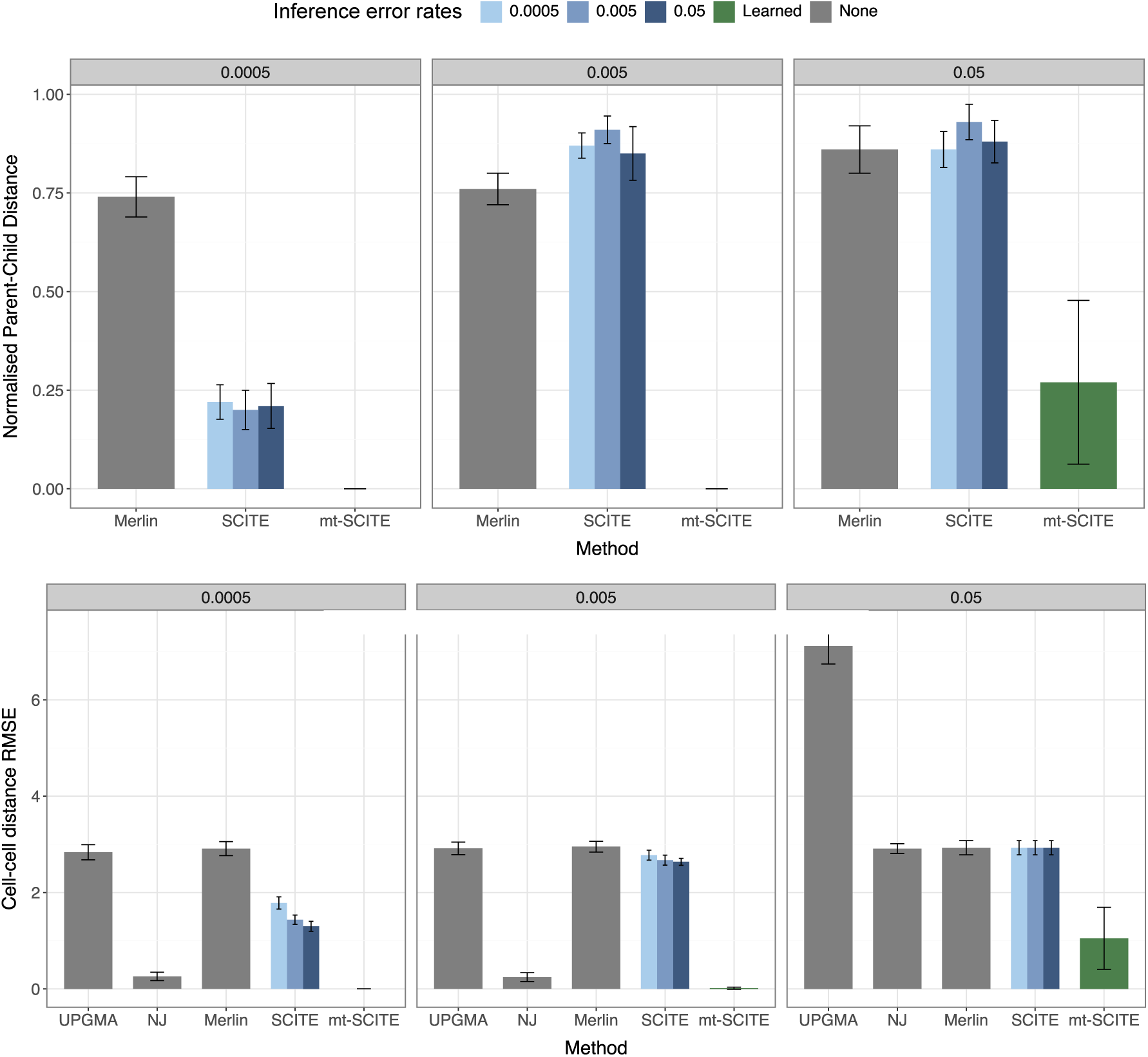
mt-SCITE outperforms existing methods in terms of mutation tree (top) and cell lineage (bottom) reconstruction on simulated data. Each panel represents input data generated with a true error rate (0.0005, 0.005, and 0.05, left to right). Bars show the normalised parent-child distance (top) and the mean square error of the cell-cell distance matrix (bottom), where lower values represent better inference accuracy. Colors denote the error rates used during inference: MERLIN, NJ and UPGMA do not account for error rate (‘None’), SCITE was evaluated for three fixed error rates (0.0005, 0.005, and 0.05), and mt-SCITE was evaluated for the learned error rate. Error bars indicate the standard deviation around the mean. Results are shown for 10 mutations, an initial mutation frequency of 10%, and 100 cells.

We first examine mutation tree inference with 10 mutations and an initial mutation frequency of 10% (Figure 2 top, Supplementary Figure 5 top), a regime in which all mutations reach high heteroplasmy frequencies allowing their detection in principle. mt-SCITE achieves near-perfect tree reconstruction across replicates with learned error rates in all conditions except the most challenging one (100 cells, error rate 0.05), where the combination of high noise and limited data prevents accurate inference. Notably, at this same error rate with 500 cells (Supplementary Figure 5), mt-SCITE recovers near-perfect trees, indicating that it effectively shares information across the additional cells to compensate for high noise. For cell lineage reconstruction (Figure 2 bottom, Supplementary Figure 5 bottom), mt-SCITE and NJ both achieve near-zero RMSE at the two lower error rates, while SCITE, MERLIN, and UPGMA have RMSE higher than 1. At the highest error rate (0.05), NJ’s accuracy plateaus regardless of cell number, while mt-SCITE continues to improve with more cells. UPGMA’s RMSE rises sharply at this error rate, indicating that it is less robust against noisier mutation calls.

When increasing the number of mutations to 50 (Supplementary Figure 9, 10), the accuracy of mutation tree reconstruction decreases across all methods, which is expected given the increased complexity of the tree search. However, as the signal to noise ratio increases - either by decreasing the error rate or by increasing the number of cells - mt-SCITE can use the additional information and achieves a normalised parent-child distance below 25% in most cases. In contrast, SCITE and MERLIN frequently exceed 50% distance indicating consistently higher reconstruction error. For cell lineage reconstruction, we observe similar patterns as for 10 mutations, namely that NJ and mt-SCITE show almost 0 RMSE for the lower error rates. At the highest error rate, mt-SCITE clearly outperforms NJ and all other methods, and this advantage becomes more pronounced as more cells are added (Supplementary Figure 10).

We further evaluated how mutations with an initial frequency of 1% affect inference (Supplementary Figures 7 and 11). At the highest error rate (0.05), the absolute performance is poor across all models, with no model achieving a mean topological distance below 0.75. This is because the initial mutation frequency (*f*_init_ =1%) is on the same order of magnitude as the error rate (5%). For high error rate and low abundance of mitochondrial mutations, mutation tree inference remains challenging. Interestingly, MERLIN’s performance decreases the most when the initial mutation frequency is decreased to 1%. This suggests that MERLIN may rely on a sufficiently high abundance of mitochondrial mutations to accurately reconstruct the mutation tree, consistent with its assumption that no clone goes extinct. In terms of cell lineage reconstruction (Supplementary Figures 8 and 12), all methods perform more similarly for the two lower error rate regimes, and distance-based methods achieve comparable or slightly lower RMSE than mt-SCITE.

We ran three sensitivity analyses to assess mt-SCITE’s robustness. First, we modelled cells sharing the same mutations but more distantly related, by increasing the variance in their heteroplasmy frequencies. Second, we tested all methods at a reduced sequencing coverage of 50 reads. Third, we introduced recurrent mutations that violate the infinite-sites assumption. mt-SCITE remained accurate under all three. Higher heteroplasmy variance did not degrade inference at low-to-moderate error rates and improved it most when the underlying signal was weak (low initial mutation frequency), since the spread of mutation signal across cells gives the method more confident observations to pool; only at the highest error rate did the added variance reduce accuracy. Performance held at low coverage, consistent with mt-SCITE explicitly accounting for the uncertainty in mutation calling that sequencing error and limited coverage introduce. Finally, accuracy was robust to mild recurrent mutation. Full results are provided in the Supplementary section “Sensitivity analyses”.

### Benchmark on experimental data

To evaluate mt-SCITE against an experimentally defined ground truth, we benchmarked it on the TF1 serial-subcloning dataset of Ludwig et al. [21] alongside SCITE, MERLIN, NJ and UPGMA (Figure 3, Supplementary Figure 13). The analysis comprised 67 samples: 65 clonally expanded subclonal populations, each derived from a single founder cell, plus two bulk samples of the founding population.

**Fig. 3:**
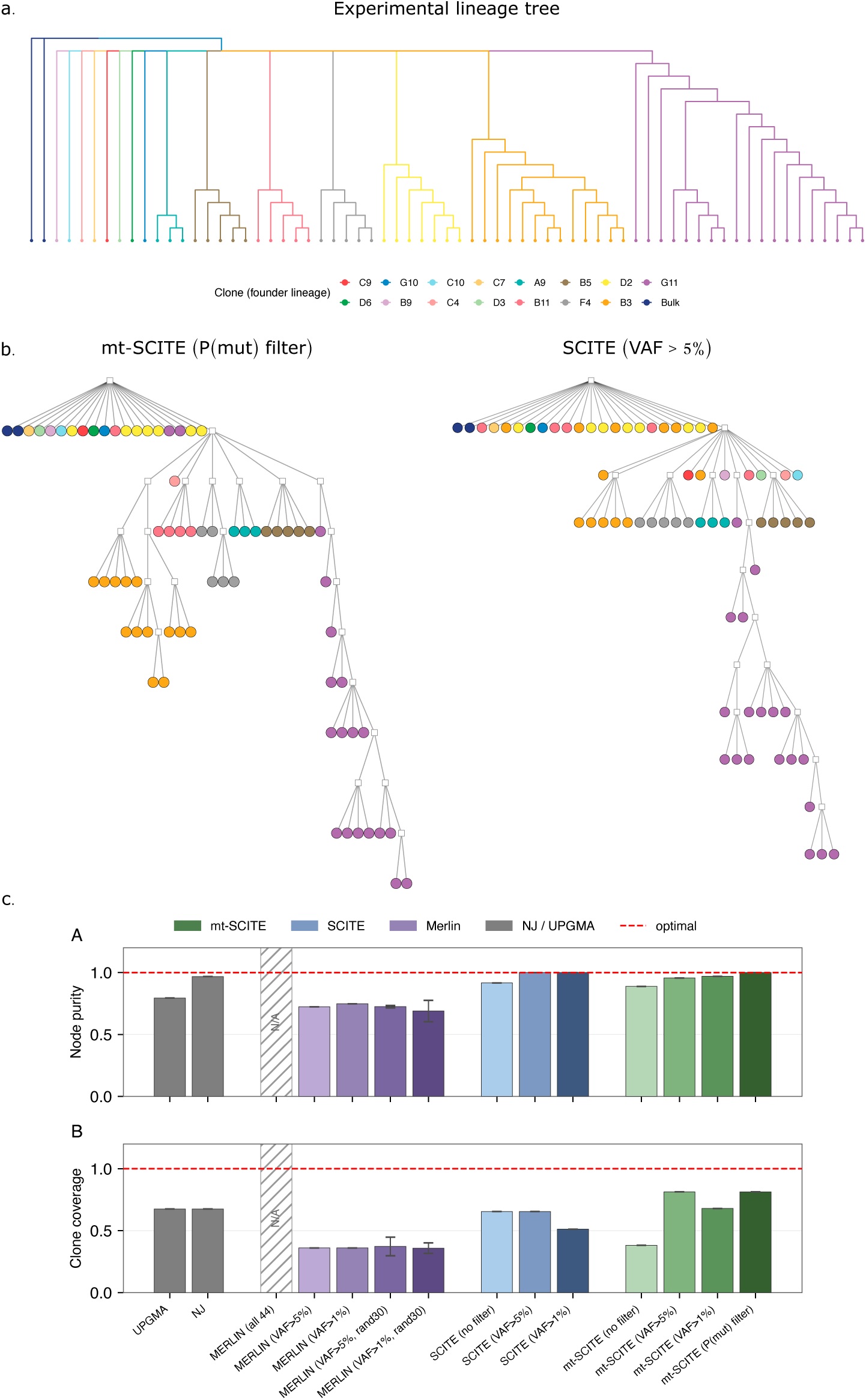
Benchmark on experimental lineage tree. (a) Ground truth cell lineage tree for the TF1 serial-subcloning experiment (adapted from Ludwig et al.). Each branching event corresponds to a culture-splitting event; colors indicate the primary founder clone from the initial split. Branch lengths are not drawn to scale. (b) Visualization of best phylogenetic trees obtained by mt-SCITE and SCITE. Mutation nodes are shown as squares and samples as circles. The samples are colored according their generating founder clone. (c) Benchmarking of all methods for 44 published variants, evaluated by node purity and clone coverage. Methods were run for variant-filtering conditions: no filter, and excluding ancestral variants detected in the bulk samples above 1% and 5% VAF, or based on the mutation probability matrix (P(mut) filter). The red dashed line indicates optimal performance.

We use the 44 variants called by the main study as input and ran all methods on the full variant set. However, mt-SCITE, SCITE and MERLIN all make the infinite sites assumption, we run them while also excluding any ancestral variants that are present at *>*1% or 5% frequency in the bulk sample. We further ran mt-SCITE using the error rate and variant learning procedure applied to the experimental datasets below (Methods). Because MERLIN is only documented to reach convergence with less than 30 variants, we also included random subsamples of the variants at 5% and 1% VAF (Supplementary Fig. 13). We score all methods in terms of node purity (the degree to which inferred subtrees contain cells from a single founder clone) and clone coverage (the fraction of each clone recovered within the most coherent subtree), which jointly assess clone recovery (Methods; Fig 3). We further evaluate the topological accuracy of the learned trees using the cell-cell distance RMSE, the Robinson-Foulds (RF) distance and the triplet distance (Supplementary Figure 13).

Across most metrics, mt-SCITE with appropriate variant filtering was the best-performing method, while SCITE and NJ also recovered the lineage well and UPGMA and MERLIN performed less consistently (Figure 3c, Supplementary Figure 13). After excluding ancestral variants detected in the bulk samples, mt-SCITE achieved the highest clone coverage (0.81) and near-perfect node purity (1.00 with its native filtering based on the mutation probability matrix; 0.96 at VAF ¿ 5%), together with the lowest Robinson–Foulds distance (0.77, compared with 0.79 for NJ and 0.82 for SCITE) and the lowest triplet distance (0.24, compared with 0.42 for SCITE and 0.45 for NJ). SCITE recovered pairwise cell–cell distances more faithfully than mt-SCITE (RMSE 0.41 versus 0.45), consistent with our simulations, in which SCITE performed best when the per-variant sequencing error rate was very low (Supplementary Fig. 12). Together, these results indicate that mt-SCITE recovered clonal structure and overall tree topology more accurately than the competing methods, with SCITE retaining a slight advantage on fine-grained pairwise distances. mt-SCITE’s accuracy depended on variant filtering: without filtering it performed worse on topology (Robinson–Foulds distance 1.00 versus 0.77 with filtering; triplet distance 0.57 versus 0.24 with filtering), reflecting the influence of ancestral variants shared across clones. Notably, its native mutation probability based variant filtering pipeline recovered strong performance (node purity 1.00, clone coverage 0.81) without requiring a manually chosen VAF threshold, supporting its use as a default in the absence of an external ground truth.

MERLIN did not run successfully on the full set of 44 variants, reporting that no perfect phylogeny existed. It completed on reduced variant sets, but retained a large optimality gap (16–18%) and performed poorly across all metrics. Restricting the input to random subsets of 30 mutations, within the operating range documented for MERLIN, narrowed the gap to approximately 6% but did not improve accuracy. Because near-optimal solutions remained inaccurate, the poor performance is unlikely to stem from incomplete optimisation alone. We instead attribute it to a model mismatch: although each sample derives from a single founder cell, the bulk sequencing of these populations pools many cells per sample, and the resulting data may fit MERLIN’s model poorly, as it strictly requires concordance between the clone and mutation trees.

### Cell lineage relationships inferred from ATAC-seq data

We evaluated the performance of mt-SCITE on clonal expansions from primary human cells (Figure 4, Supplementary Figures 2, 23). The clonal expansions were derived from single memory CD8+ T-cells isolated from a human donor vaccinated against Yellow Fever Virus (YFV) (Supplementary Figure 1) [36]. Each *in vitro* expanded clone (*n* = 6) was split into two replicates, and the resulting 12 bulk samples were subsequently subjected to ATAC-seq (Figure 4a).

**Fig. 4:**
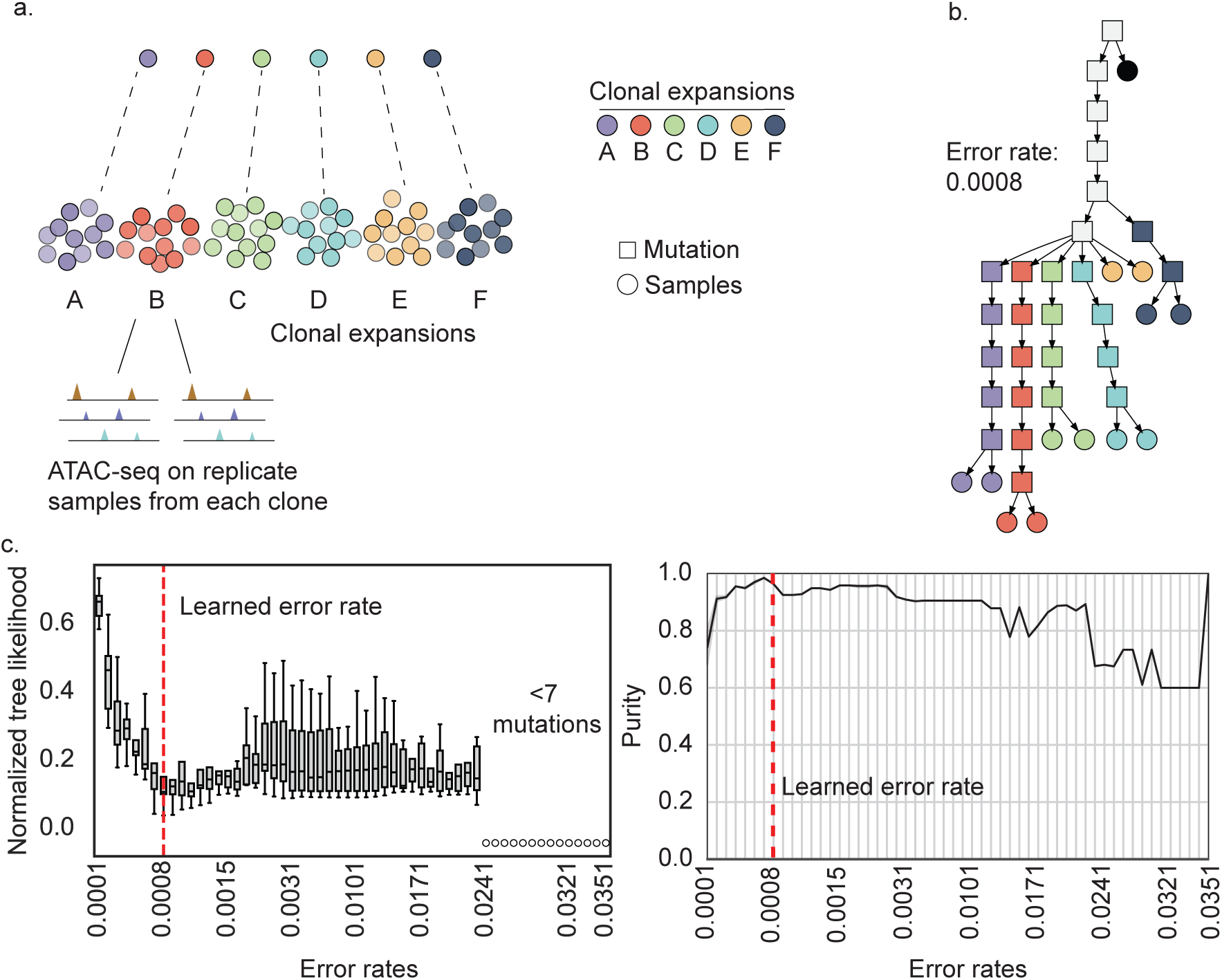
Tree inference from clonal CD8+ memory T-cell expansions subjected to ATAC-seq. (a) Clonal expansions from single CD8+ memory T-cells are split into replicates and subjected to ATAC-seq. (b) Visualization of a phylogenetic tree obtained by mt-SCITE. Mutation nodes are represented by squares and samples are represented by circles. The samples are colored according to the clonal expansion they originate from. Pure nodes are labeled in the same color as the clonal expansion replicates attached to them. (c) The error rate of the sequencing experiment was learned using 3-fold cross-validation with 3 repetitions. The boxplots display the normalised tree likelihood across all folds and repetitions per error rate. (d) The average tree purity across the evaluated error rates, excluding mutation nodes attached to all samples.

From these data, we used mt-SCITE to compute mutation probability matrices for a range of error rates and then filtered out mutations that were unlikely to be part of the tree (Methods). As expected, the number of detected mutations increased as the error rate decreased (Supplementary Figure 2b), because low error rates make it more likely that even small non-reference read counts represent true mutations.

Next, we learned the error rate of the sequencing experiment by performing 3-fold cross validation with 3 repetitions (Methods). For the analysed ATAC-seq dataset, the learned error rate was 0.0008 (Figure 4c). Additionally, we used mt-SCITE for phylogenetic inference on the full dataset for every error rate and assessed the reported trees.

We evaluate the inferred trees using tree purity and clone coverage, which capture aspects similar to precision and recall, respectively, of how well the tree recovers clonal structure. Tree purity measures, for the samples grouped under each internal node, the average fraction that come from a single dominant clone. Clone coverage measures the complementary quantity: for each clone, the largest fraction of its members that the tree gathers under a single, clonally pure node.

We found that the learned error rate falls within a range of error rates that exhibit similarly high purity, with 96% average purity for the learned error rate. Further, the ML tree found at this error rate showed separated clones (A-F), with each clone forming a distinct subtree and each set of replicates per clone attaching to the same mutation node. mt-SCITE maintains high purity over a broader range of error rates, demonstrating the robustness of the approach. Taken together, these results demonstrate the utility of mt-SCITE in identifying samples with common ancestry.

### Cell lineage relationships inferred from single-cell RNA sequencing of expanded T-cells

Next, we resolved cell lineage relationships *in vivo*. We analysed a scRNA-seq dataset obtained from T-cells isolated from a healthy human donor vaccinated against YFV [35, 37]. The vaccination triggers the activation of a small subset of näıve T-cells which clonally expand and enter the circulation. These cells can subsequently be longitudinally collected directly from blood (Supplementary Figure 2).

T-cells represent a useful system for evaluating the performance in identifying clones in real data, because clonally related T-cells share a unique genomic sequence in nuclear DNA, the TCR. Since the TCR is expressed, it can be detected in scRNA-seq data and be used as ground truth for clonality. To evaluate the performance in identifying clonal populations of human cells, we selected 72 cells belonging to six T-cell clones with support for at least one mitochondrial mutation in the cells sharing the same TCR (Figure 5, Supplementary Figures 3, 24). These cells were sampled on days 15, 30, and 593 post-vaccination, representing samples of the acute phase (days 15 and 30) and late memory (day 593) of the immune response [35]. In addition to the single T-cells, we included an unamplified bulk sample comprising PBMCs isolated from the same donor, which was subjected to WGS as this allows us to distinguish germline mutations from somatic mitochondrial mutations.

**Fig. 5:**
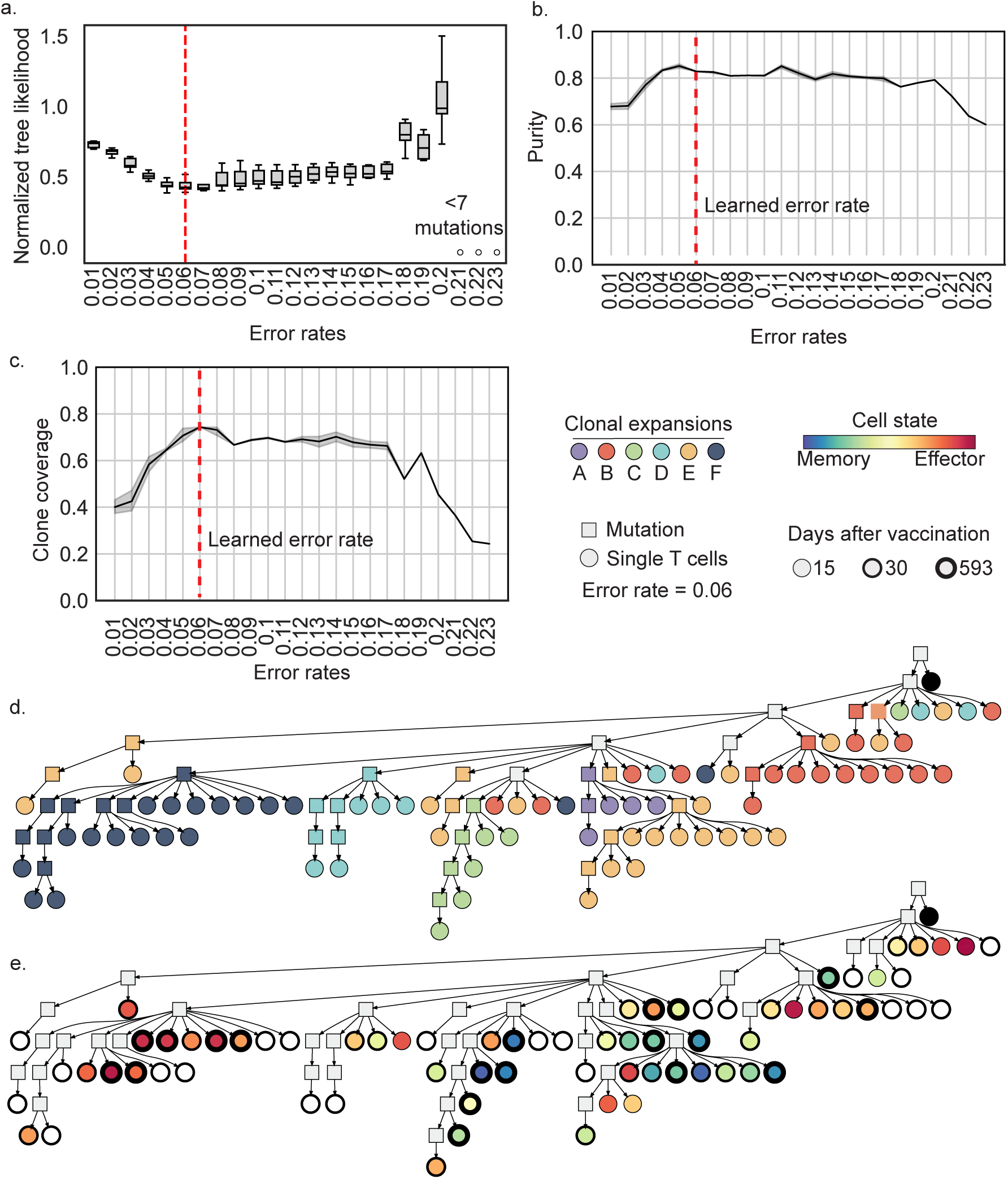
Tree inference from scRNA-seq data comprising *in vivo* expanded CD8+ T-cell clones. (a) The error rate of the sequencing experiment was learned using 3-fold cross-validation with 3 repetitions. The boxplots display the normalised tree likelihood across all folds and repetitions per error rate. (b) Average tree purity across the evaluated error rates, excluding the root and any node from which all cells descend. (c) Average clone coverage across the evaluated error rates. (d) A phylogenetic tree comprising *in vivo* expanded CD8+ T-cell clones. The cells (circles) are colored by the TCR clone they belong to. Mutation nodes (squares) are assigned the color of a TCR clone if they are pure and grey if cells from multiple clones are attached to it. (e) The phylogenetic tree from (d) annotated by cell type classification where colors indicate cell state and edge width represents the collection timepoint. White represents cells with an undetermined classification.

We used mt-SCITE to compute the mitochondrial mutation probabilities and to build trees of the 72 single T-cells and the unamplified bulk sample. We applied a 3-fold cross-validation approach with 3 repetitions to estimate the sequencing error rate, which was determined to be 0.06 (Figure 5a). In accordance with the ATAC-seq dataset, the number of detected mutations increased with decreasing error rates (Supplementary Figure 3b).

The bulk sample was attached directly to the root node and thus did not share somatic mitochondrial mutations with any of the single cells. This is expected because the bulk sample consists of millions of cells, and signals from somatic mitochondrial mutations that occur in only a small subset of those cells become diluted and effectively obscured in the overall data. We expect that clonally related cells, defined as sharing the same TCR, will be attached to the same mutation nodes in the tree, reflecting the inheritance patterns of mitochondrial mutations. However, given errors in the data, the high mutation rate in mtDNA and genetic drift, the data may not perfectly support the clonal structures. The performance of mt-SCITE to identify clonal populations was assessed by analyzing the purity of individual mutation nodes across the trees. mt-SCITE achieved 83% average tree purity for the learned error rate 0.06 (Figure 5b). In addition, we measured the clone coverage in pure nodes. We found that mt-SCITE achieved 74% average clone coverage in the T-cell dataset (Figure 5c).

We visualised the distribution of clonal T-cell populations across the tree by labeling cells sharing the same TCR (Figure 5d). Apart from clone E, cells from the same founder clone cluster together in the phylogeny. Since the mutations are called directly in the scRNA-seq data, we could use the same data to superimpose cell type classifications on the tree to visualize the genealogical relationships between distinct T-cell subsets. This revealed clones where the majority of cells had either a memory (clones A, C) or effector phenotype (clone F), as well as clones with both memory and effector CD8+ T-cells (clones B, D, E). This finding was further confirmed by visualizing effector and memory gene expression heterogeneity within the tree structure (Figure 5e, Supplementary Figure 4) [35, 37]. Taken together, mt-SCITE allowed us to robustly identify clonal cell populations *in vivo* and revealed the evolution of CD8+ T-cell clones after vaccination against YFV.

### mt-SCITE infers cell lineage relationships from large scale single-cell RNA sequencing data

To demonstrate scalability, we applied mt-SCITE to the MAESTER dataset [22], comprising 1,397 cells and 26 mutations. Parallelised across 10 cores, the full analysis, including learning the sequencing error rate via 3-fold cross-validation with 3 repetitions and inferring the trees, completed in 1 h 5 min. This efficiency is expected from how the likelihood scales. Each cell’s attachment is marginalised independently over the n + 1 possible positions in the tree, so the per-MCMC-step cost grows only linearly in the number of cells. The bottleneck is therefore not cell number but tree-space exploration, which grows super-exponentially in the number of mutations *n*. The 3-fold cross-validation procedure estimated an error rate of approximately 0.0089 (Fig. 6a). This is consistent with the quality filters applied in the original study [22], which retained variants with mean base quality *>*Q30 and applied further filtering steps based on coverage and required VAF (see methods for details). A Phred score of Q30 corresponds to a per-base error probability of 10^−3^, but since only the mean is constrained, individual bases at lower quality can contribute higher error rates. In addition, the learned rate captures not only sequencing errors but also PCR amplification errors and residual noise after UMI-based consensus calling, collectively placing the realised error rate plausibly above 10^−3^. Our estimate of 0.0089 remains below the 1% single-cell VAF threshold used in the original study to call a cell positive for a variant, confirming that the signal for a variant is separable from technical noise.

**Fig. 6:**
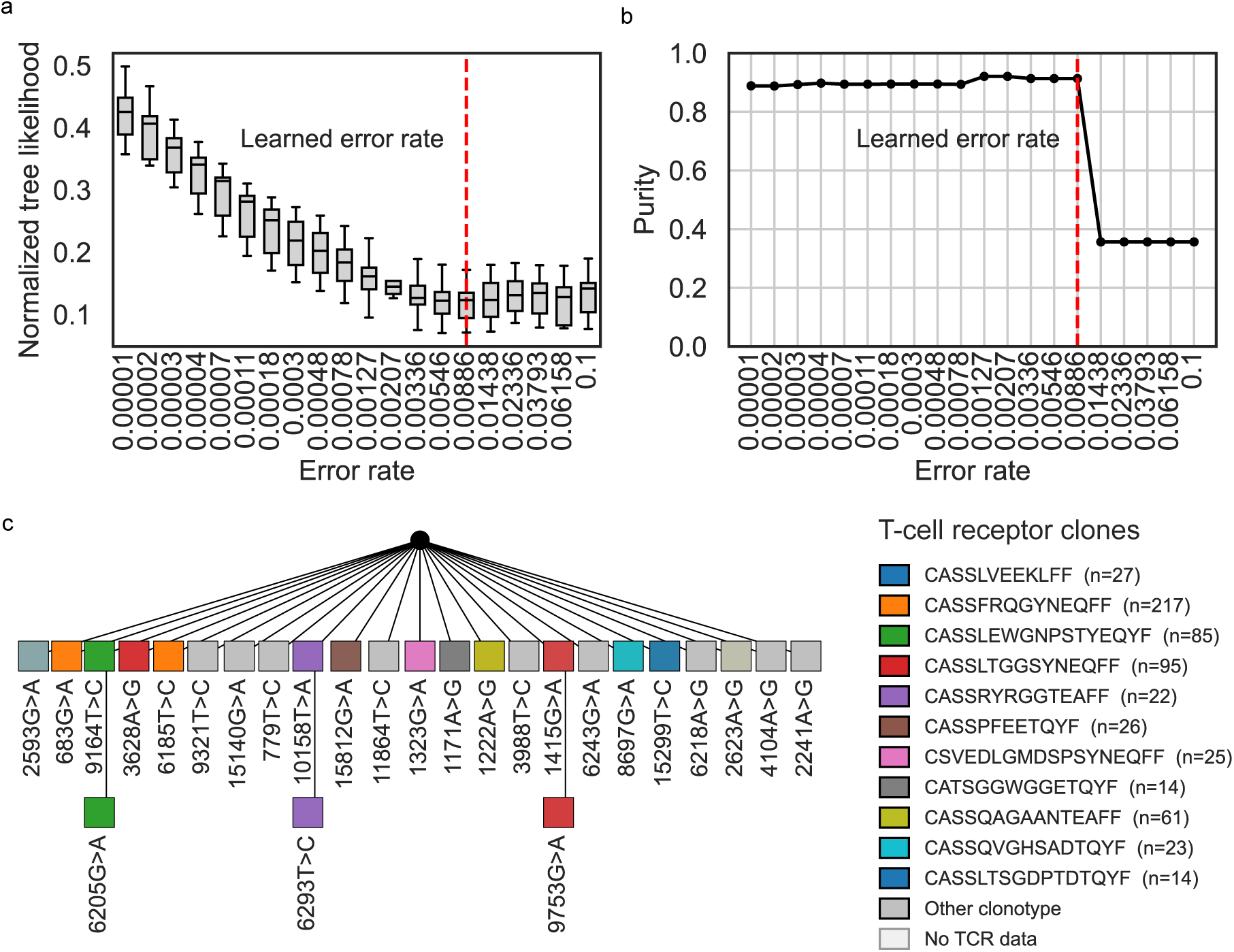
Tree inference from MAESTER scRNA-seq data of a bone marrow aspirate (1397 cells and 26 mutations). (a) The sequencing error rate was learned via 3-fold cross-validation with three repetitions. Boxplots display the normalised tree likelihood across all folds and repetitions for each error rate. (b) Mean tree purity across the evaluated error rates. (c) Visualization of a phylogenetic tree of mitochondrial mutations (squares). Cell node attachments are omitted from the visualisation to avoid crowding. To validate the tree structure, we colour the mutation nodes by the T-cell receptor (TCR) clones of their descendant cells. Cells lacking TCR annotations are transparent and thus do not contribute to the mixture of colours.

The MAESTER dataset profiles the T-cell receptor (TCR) using TREK-seq, which sequences the TRA and TRB variable regions from the 10x 3’ scRNA-seq cDNA, providing an orthogonal lineage annotation alongside the mitochondrial variants for 609 of the 1397 cells. We use the TRB region as an independent lineage readout and compute the purity of TCR clone assignments for the descendant cells of each mutation node, as in the previous section. The inferred tree achieves a mean purity of 91%, indicating that mitochondrial mutations carry substantial information for reconstructing lineage relationships (Fig. 6b).

Examining the tree structure, we observe two informative patterns. First, we observe pairs of mutations that independently mark subsets of cells within the same TCR clonotype. For example, 683G*>*A and 6185T*>*C each define separate top-level branches of the tree, yet both consist predominantly of cells from the CASSFRQGYNEQFF clone (Fig. 6c). Because these mutations lie on independent lineages but are confined to a single TCR clonotype, they must have been acquired after TCR rearrangement, and together they reveal subclonal structure within a mature T-cell clone that would be invisible to TCR sequencing alone. This is consistent with observations in Miller et al. [22], who reported concordance between mtDNA clones and TRB clonotypes including cases where an mtDNA clone subdivides a single TCR clonotype. Our phylogenetic reconstruction places these relationships in an explicit tree, adding the order and branching structure of the underlying mutations. Second, the mutation 2593G*>*A is shared across multiple TCR clonotypes and, as reported by Miller et al. [22], is also detected in non-T-cell lineages; together this places its origin in a multipotent hematopoietic progenitor, upstream of TCR rearrangement.

## Discussion

The rapid development of DNA sequencing technologies and customised data analysis tools has recently enabled cell lineage tracing in humans, taking advantage of the inheritance patterns of somatic variation in individual cells. Utilizing mitochondrial mutations for lineage tracing is a particularly interesting approach, since these can be detected in ATAC-seq and scRNA-seq data, which opens the possibility for combined analysis of cell lineage relationships and cell states. However, inferring cell lineage relationships from shared mitochondrial mutations is challenging because of the complex mutation inheritance dynamics in mitochondria [15] and multiple types of error in single-cell data, such as false positive and false negative mutations.

We have presented mt-SCITE, a method for inferring cell lineage relationships based on shared mitochondrial mutations. In our workflow, we determine the probability of whether a mutation is present in a particular locus and sample. These mutation probabilities are used for computing the maximum likelihood evolutionary history of the samples. The framework allows us to simultaneously account for errors occurring during the sequencing experiment and to robustly infer cell phylogenies as demonstrated on simulated and real data. Our tree inference approach represents an advantage over currently used clustering methods since it allows us to explicitly model the genealogy of dividing cells and accounts for sequencing errors. Further, mt-SCITE outperforms a recently introduced method designed for tree inference from mitochondrial mutation data.

We validated mt-SCITE on a dataset of *in vitro* expanded T-cell clones following vaccination of a healthy donor with the Yellow Fever Virus and estimated a cell lineage tree with high purity and clone coverage, measures of clonal recovery. We further benchmarked mt-SCITE against an experimentally defined ground truth using the TF1 serial-subcloning experiment of Ludwig et al. [21], in which the culture-splitting pedigree provides a known lineage topology. Here, mt-SCITE recovered clonal structure with high fidelity, achieving near-perfect node purity and the highest clone coverage among the methods compared; however, recovering the exact branching hierarchy proved more difficult. Topological accuracy was only moderate (Robinson-Foulds distance 0.77, triplet distance 0.24), albeit again the best of the methods tested. While complete phylogeny inference remains challenging at current sequencing depths and number of variants detected, mt-SCITE is capable of robustly identifying clonally related populations together with meaningful substructure within them.

We further applied mt-SCITE to the question of T-cell expansion following vaccination, with repeatedly donated T-cells that were profiled using single-cell RNA sequencing [35]. In the original study, the authors tracked T-cell differentiation and clonal expansions primarily through TCR sequencing and single-cell transcriptomics. Here, we integrate an additional layer of lineage information through mitochondrial mutations using mt-SCITE, thereby enhancing resolution of subclonal relationships beyond TCR-defined clones. This allows us to dissect the genealogical relationships even among T-cells that share the same TCR but exhibit distinct phenotypic fates, refining our understanding of how clonal architecture evolves during the CD8+ T-cell response to vaccination (Figure 5d). Overall, our findings support published reports demonstrating that somatic mutations are present in mtDNA of T-cells and can be used to distinguish clonal cell populations in humans [19–23, 26, 42].

The reliability of mitochondrial mutations as lineage markers depends on the biological context. Wang et al. [43] showed in simulations that mitochondrial variants distinguish lineages reliably only under strong, recent clonal expansion; otherwise the called variants tend to be pre-existing founder heteroplasmies rather than de novo mutations acquired along the lineage. Campbell et al. [44] reached a consistent conclusion using single-cell whole-genome lineages as ground truth. Both delimit the regime in which mitochondrial mutations carry phylogenetic signal: strong clonal expansion over timescales which are short relative to heteroplasmy drift. Our datasets fall squarely within this regime, especially the T-cell expansion following Yellow Fever Virus infection, so mt-SCITE’s good performance is consistent with these findings. We also saw that removing pre-existing founder variants helped improve lineage reconstruction, most clearly on the TF1 dataset, where excluding ancestral variants improved topological accuracy. Within the clonal expansion regime, mt-SCITE not only recovers clones with high purity but also resolves structure below the clonal level, demonstrating that mitochondrial phylogenetics provides useful additional information when the context supplies the signal.

While our findings demonstrate the utility of mt-SCITE for lineage reconstruction, several avenues for future research remain to further refine and expand its applications. A key challenge is the limited understanding of how mitochondrial mutations evolve within a cell and are inherited by daughter cells [15, 43]. This compelled us to make simplifying assumptions when designing our tree inference model. For instance, our model assumes that descendant cells will still retain some fraction of the mutation from their parent cell, without explicitly modelling the expected allele fraction. Additionally, the prior probability of a mutation is currently treated as constant across the mitochondrial genome. In reality, mutation rates vary across mtDNA loci, and the mutational spectrum varies in magnitude and composition between tissues and with age [45]. Building more complex models that capture more of the known biology and comparing them to mt-SCITE in terms of accuracy and scalability is an important direction for future work.

Another consideration is the infinite sites assumption, which is shared by SCITE and Merlin but not Neighbour joining (NJ) and UPGMA. According to this assumption, every mutation arises only once in the history of the cells. However, given the high mutation rate in mtDNA, the same mutation may appear independently in unrelated cells which could lead to biases in tree reconstruction. Despite this possibility, it remains unclear how frequently such homoplasy occurs, and over what experimental timescales it becomes a prevalent problem in the data. Addressing homoplasy would require incorporating the possibility of recurrent mutations into the inference framework, which would significantly increase the complexity of the tree search [46]. Nevertheless, our sensitivity analyses show that mt-SCITE tolerates modest levels of recurrent mutation, indicating that limited homoplasy has little impact on tree reconstruction. Notably, the infinite-sites-free methods NJ and UPGMA do not beat mt-SCITE under homoplasy: on cell-cell distance reconstruction mt-SCITE remains at least as accurate as NJ and more accurate than UPGMA across all conditions tested.

Scaling mitochondrial phylogenetics toward fully cell-resolved trees will require both a deeper understanding of mitochondrial mutation dynamics and datasets that carry more phylogenetic signal. Simulation-based studies may clarify how different mutation and population regimes shape the reliability of mtDNA as a lineage marker [43], and integrated readouts of mitochondrial variation alongside genetic barcoding [47] could provide ground-truth datasets for studying how mitochondrial mutation frequencies evolve along the cell lineage tree. A complementary route is simply more informative variants, a number that is rising rapidly. Deep mitochondrial profiling approaches such as ReDeeM [47] use single-molecule consensus error correction to call an order of magnitude more somatic variants per cell, including the low-heteroplasmy mutations that they argue carry much of the fine-scale lineage signal. Because mt-SCITE operates on per-base read counts and models sequencing error explicitly rather than relying on hard genotype calls, it is well suited to the more permissive variant sets such methods produce, and can down-weight weakly supported calls rather than treating every called variant as equally certain. Phylogenetic signal could also be increased by combining complementary sources of lineage information. Coupling the likelihood of mitochondrial variants with nuclear variants called from the same single-cell RNA-seq reads, for example, is a worthwhile direction. Overall, as the understanding of mitochondrial dynamics deepens and confident variant sets grow, computational models such as mt-SCITE will recover increasingly fine-grained cell lineage relationships in humans.

## Methods

### mt-SCITE input data

mt-SCITE takes two inputs: (i) a list of per-sample read-count matrices covering every informative position in the mitochondrial genome, and (ii) a CSV file giving the reference nucleotide at each of those positions. A sample is the unit attached to the inferred tree: a single cell for single-cell assays (scRNA-seq, scATAC-seq; Figs. 5, 6) or a bulk-sequenced clonal population for the bulk ATAC-seq data (Figs. 3, 4). Each per-sample read-count matrix has entries *v_ij_*, where *i* indexes mitochondrial positions and *j* ∈ {*A, C, G, T* } indexes the four nucleotides, so that *v_ij_* is the read count supporting nucleotide *j* at site *i* in that sample. From these, mt-SCITE computes per-sample, per-variant mutation probability matrices (see Section mt-SCITE mutation probabilities) that are passed to the mutation tree inference (see Section mt-SCITE tree inference).

This input format is agnostic to the upstream assay and variant-calling pipeline. mt-SCITE is therefore directly applicable to data from many modalities that yield mitochondrial read counts. In this manuscript, we demonstrate its use on MAESTER and related 3’-biased scRNAseq protocols, scATAC-seq or bulk ATAC-seq; other modalities such as mtscATAC-seq [20] can be used without modification. Coverage uniformity and per-position depth vary between assays and affect the accuracy of the mutation probability estimates, and therefore of the downstream tree inference, but require no methodological changes to mt-SCITE itself.

### Model overview and assumptions

mt-SCITE comprises three components. The first is a within-sample observation model (Section mt-SCITE mutation probabilities): for each site and sample it computes the posterior probability that a mutation is present, given the per-nucleotide read counts, the total coverage, and a given sequencing error rate, under a fixed prior probability of mutation per site. This yields a mutation probability matrix *A* of soft, probabilistic genotype calls rather than hard present/absent assignments.

The second is a tree likelihood (Section mt-SCITE tree inference): given a mutation tree and an attachment of samples to its nodes, the likelihood of each sample is the product, over all sites, of the probability that the sample carries the mutations on the path from the root to its attachment point and does not carry the remaining mutations, read directly from the entries of *A*. The likelihood of a tree is the product of these per-sample terms, and inference searches for the tree and attachment that maximise it.

The third learns the sequencing error rate (Section mt-SCITE error rate learning). Because the true rate is generally unknown, mt-SCITE evaluates tree inference across a grid of plausible error rates and, using cross-validation, selects the rate whose inferred tree best generalises to held-out samples.

Taken together, from per-site, per-sample read counts mt-SCITE returns a maximum-likelihood tree and an estimated sequencing error rate.

### mt-SCITE mutation probabilities

The workflow of mt-SCITE begins with determining the probability of whether any mutation is present at a particular site in a particular sample. For each site *i* and sample *j*, let the read counts supporting each nucleotide be denoted by 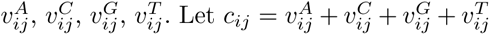 be the total coverage.

The aim is to generate a mutation probability matrix **A** of size *L* × *N* (Figure 1), where *N* is the number of samples (cells or bulk populations) and *L* is the number of sites (or mutations). Each entry *A_ij_*is the probability that sample *j* carries a mutation in site *i*.

We define each site *i* in sample *j* as being in one of four possible states: the unaltered reference state (*R*) or one of three alternative nucleotide states (*X, Y, Z*). We assume that mutations can only cause a transition from *R* to one of the alternative states.

To compute *A_ij_*, we sum over the probabilities of a mutation into any of the non-reference nucleotides, X, Y or Z, given the observed read counts, total sequencing coverage and the error rate *ρ*,

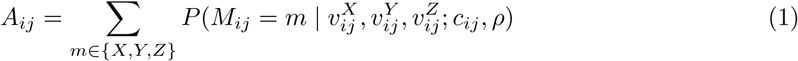

Note that *M_ij_* can take any of the four values *R, X, Y, Z*. However, the sum defining *A_ij_* excludes the reference state (*m* = *R*), so *A_ij_* is the total probability that the site is in one of the mutated states *X, Y, Z*.

We express the posterior probability of the mutation state *M_ij_* = *m* given the observed read counts and error rate using Bayes Theorem:

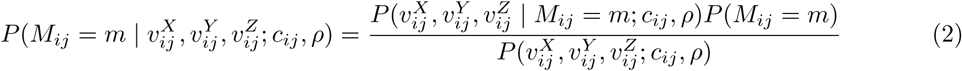

where 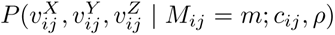 is the likelihood, the probability of observing the read counts given that the true mutation state is *m*; *P* (*M_ij_* = *m*) is the prior probability of the site being in state *m*; and 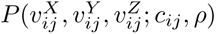 is the evidence, which normalises the posterior. In the following, we detail how each of these terms is computed.

For computing the prior probability, we assume that observing each alternative nucleotide is equally likely. Based on the findings from previous investigations [36], we expect the number of mutations to be on the order of 10 across the 16.6kb mitochondrial genome, corresponding to a prior probability of mutation 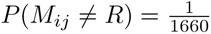. Thus, the prior probability of each alternative nucleotide is:

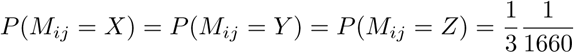

The prior probability of a site remaining in the reference state is 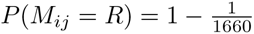

The evidence, or the probability of observing the read counts 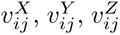 conditional on the coverage and error rate, can be expressed as the marginal probability obtained by summing over all possible states *M_ij_* ∈ {*X, Y, Z, R*},

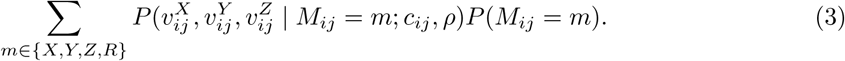

To compute the likelihood, we distinguish between two cases:

1. A mutation is present in one of the alternative nucleotides (*M_ij_* ∈ {*X, Y, Z*}).
2. No mutation is present, meaning the true nucleotide corresponds to the reference state (*M_ij_* = *R*).

### Case 1: A mutation is present

Let there be a mutation in nucleotide X. Using the chain rule, we can express the likelihood as

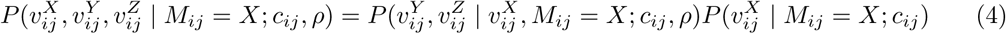

where the first term represents the probability of observing read counts for nucleotides Y and Z, given that we already observed 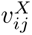 read counts for X and that X is the true state. The second term represents the probability of observing 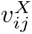 reads for X given that X is the true state.

Since reads supporting Y and Z arise only through sequencing error, and sequencing errors are assumed to be distributed uniformly among the two erroneous nucleotides and independently across sites, each erroneous read occurs with probability *ρ/*2. Thus, the first term follows a multinomial distribution,

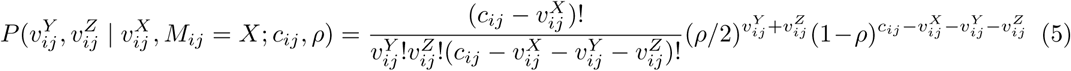

For the second term, 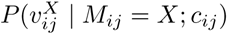, we approximate the probability of observing 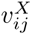 reads for X, given that X is the true state. Since we assume no knowledge of the underlying heteroplasmy level we approximate this term with a uniform distribution 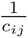 over *c_ij_* possible counts 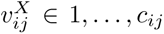 and include an alternative derivation in the Supplement (“Variant read count approximation”).

### Case 2: No mutation is present

If no mutation is present, meaning the true nucleotide is the reference state *R*, any observed reads for X, Y or Z arise solely from sequencing errors. Under this assumption, each read has an independent probability *ρ* of being miscalled due to sequencing error. Since there are three possible alternative nucleotides (X, Y, Z), and errors are assumed to be equally likely across them, the probability of a reference nucleotide being incorrectly identified as any specific alternative is *ρ/*3. Thus, the probability of observing read counts 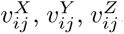, follows the multinomial distribution

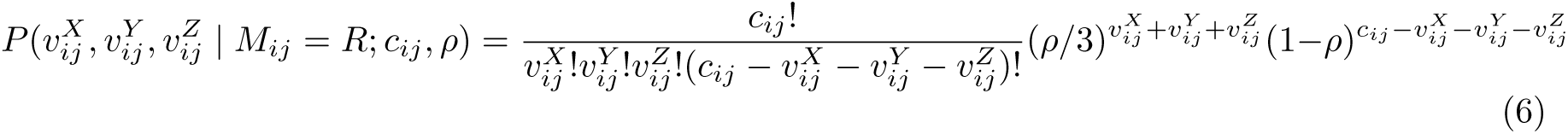

### mt-SCITE tree inference

Cell lineage relationships were inferred by mt-SCITE, a new statistical model and stochastic search and sampling algorithm for phylogenetic inference taking mitochondrial mutation probabilities in individual cells as input. mt-SCITE adapts [38], which works with binarised mutation presence/absence data, to handle instead mutation frequencies as present in mitochondrial data. The output of mt-SCITE consists of a maximum likelihood mutation tree, which represents the estimated genealogical relationships between the individual cells in the dataset. In this tree, each node *i* corresponds to a mutation in a site *i*. That is, the index *i* of a node in the mutation tree is identical to the index of the corresponding mutated site in the mutation probability matrix **A**. Further, every cell *j* is attached to its attachment node *σ_j_*. The mutational profile of each cell is obtained by accumulating the mutations from the root of the tree to the node the cell is attached to. The model uses the mutation probability matrix **A** as input data to define the likelihood of the data for a tree structure *T* and cell to node attachment vector ***σ***. The likelihood contribution of each cell is determined by the product of the probabilities of each mutation along the tree that leads to the attachment point of the cell. Let anc(*T*)*_j_* denote the set of nodes in the path from the root to and including the node *σ_j_*, which is the node that cell *j* attaches to. Then, the likelihood is given by

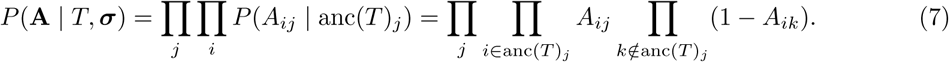

The algorithm uses a search-and-score procedure to find the tree *T* ^∗^ that maximizes the probability of the data over all trees and attachment vectors:

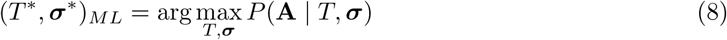

mt-SCITE starts by initializing a random mutation tree structure on the input mutations, and proceeds by iteratively perturbing the mutation tree with local moves: (i) pruning branches and reattachment to other locations in the tree, (ii) swapping node labels, and (iii) swapping subtrees. Note that (i) is the equivalent of the subtree pruning and regraft (SPR) move on a mutation tree. These moves are described in detail in [38].

mt-SCITE outputs all the trees with the same optimal score that were encountered during the search procedure. Depending on the error rate used to compute the posterior mutation probabilities that are used as input for the tree inference, the number of trees with the same optimal score may differ. In principle, the higher the prior error rate, the lower the posterior mutation probabilities, and thus a larger number of different trees with the same ability to explain the data is expected. In contrast, if the prior error rate is lower, mutation calls are more confident and the tree space that explains the data accurately is more restricted.

### mt-SCITE error rate learning

We have introduced above how to estimate the mutation probability matrix and reconstruct the tree for a given error rate. However, in most cases, the true error rate is unknown. For simplicity, we learn trees via MCMC over a range of candidate error rates and then select the one that best fits the data. Specifically, we use a *k*-fold cross-validation approach, in which we partition the mutation probability matrix by cells into *k* folds. For each error rate, we estimate a maximum likelihood tree using mt-SCITE on the training set. We then take this maximum likelihood (ML) tree and evaluate its log-likelihood on the test set, which we define as log L(T_train_ | A_test_). We further normalise this likelihood against the log-likelihood an uninformative star tree would achieve on the test set, correspondingly defined as log L(T_star_ | A_test_). We choose the error rate that minimizes this ratio. This yields the following criterion:

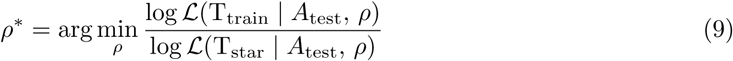

Minimizing this ratio ensures that the inferred tree provides a better fit to the data than the baseline star tree, thereby avoiding the selection of excessively high error rates. That is because at high error rates, no mutations are supported and every tree is equally capable of explaining the data. By normalizing against the log-likelihood of a star tree, we can effectively reject error rates in this regime. Because the star tree’s log-likelihood is negative, we minimize the ratio rather than maximize it.

We also opted against a direct likelihood ratio test between the training tree and the star tree since the ratio directly accounts for different potential sizes of data depending on the mutation filtering, and hence different potential scales of the likelihoods.

### Performance evaluation

We assess the reconstructed tree against the ground-truth clone labels (the T-cell receptor labels) using two complementary metrics, tree purity and clone coverage, capturing notions of precision and recall of clone recovery.

### Tree purity

Let *S* be the set of all samples and *S_g_*(*c*) ⊆ *S* the samples belonging to **g**round-truth clone *c*. For a node *n* in the inferred tree, let *S_d_*(*n*) = { *s* ∈ *S* | *s* is a descendant of *n* } be the set of samples attached at or below *n*. We define the fraction of the descendants of *n* belonging to clone *c* as

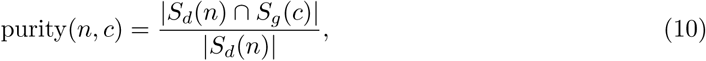

and the *node purity* as the fraction belonging to its dominant clone, node purity(*n*) = max

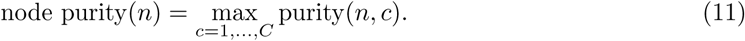

Node purity is a precision-like measure: it equals 1 when all descendants of *n* belong to the same clone, and takes its minimum value 1*/*|*S_d_*(*n*)| when every descendant belongs to a different clone.

To summarise a whole tree *T*, we average node purity over the set of informative internal nodes

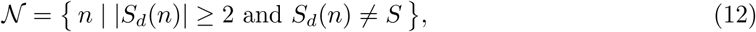

excluding single-descendant nodes, which are trivially pure, and the node from which all samples descend, which cannot be pure whenever more than one clone is present. The *tree purity* is

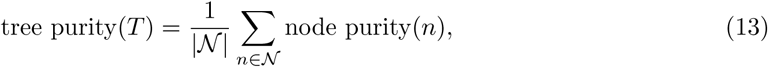

which equals 1 when every informative node is clonally homogeneous.

**Clone coverage** is the complementary, recall-like measure: for each clone, the largest fraction of its members that the tree gathers under a single, fully pure node. Defining the set of pure nodes

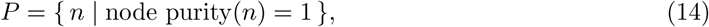

the clone coverage is

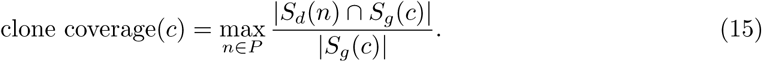

A clone with coverage 1 has all of its samples isolated as descendants of a single node carrying no other clone. As a tree-level summary we report the mean clone coverage across clones; we additionally report the minimum across clones,

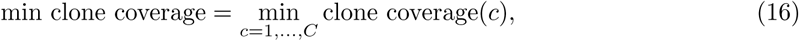

as a conservative worst-clone score.

### Method comparisons

We compared mt-SCITE to SCITE, MERLIN, Neighbour joining (NJ) and the Unweighted Pair Group Method with Arithmetic Mean (UPGMA), on mitochondrial mutation trees (SCITE and MERLIN) and cell lineage trees (all) using simulated data with known ground truth trees and on an experimental dataset with ground truth cell lineage tree from a cell splitting experiment in culture [21] (Fig 1c).

### Data simulation

To simulate mutation trees, we generated random trees with *L* mutations across *L* + 1 nodes. The starting node with index 0 represents the wild-type state without mutations, and each additional node *i* introduces a new mutation *i*. Each node *i* has an associated frequency vector *f_i_*of length *L*, where *f_ik_* represents the frequency of mutation *k* at node *i*. The frequency vector for the starting node is initialised as *f*_0_ = [0*, . . .,* 0].

To introduce a new (child) mutation *c*, we randomly select a parent node, *p*, from the existing tree and attach the child node to it, equivalent to growing a random network with uniform attachment [48]. Note that we number the nodes by the introduction order, meaning the starting wildtype node is 0, the first introduced mutation node is 1, et cetera. We next sample the mutation frequencies at the child node and distinguish between the newly introduced mutation *c* and all potential ancestral mutations *k* ∈ {1*, …, c* − 1}. We draw the frequency of the newly introduced mutation from a Beta distribution. We use the Beta distribution because its support is exactly [0, 1], making it a natural choice for an allele frequency and we adopt the mode/concentration re-parameterization [49] which lets us fix the central tendency to a biologically motivated value (the mode *o* = *f*_init_) while controlling the spread through the concentration parameter *κ*:

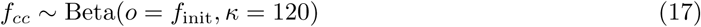

In our simulations, *f*_init_ is set to 1% and 10% to reflect biologically plausible starting frequencies. A frequency of 1% represents a single mutation in a pool of 100 – 1000 mitochondrial genomes [21], while 10% corresponds to a scenario where we only keep track of a mutation once it has already reached a high enough frequency to be detected at the end of the experiment.

We chose *κ* = 120 as it produces a Beta distribution with a reasonable spread around the mode (Supplementary Figure 22). Specifically, for mode 0.01 the 5% quantile is 0.004 and the 95% quantile is 0.04, allowing minor variation around the mode. This ensures that the child mutation frequency is on the same order of magnitude as the parent mutation frequency, preserving a high phylogenetic signal in the mutation frequencies.

For all potential ancestral mutations *k* ∈ {1*, …, c* − 1}, we resample their frequencies based on the parent node frequencies, to mimic the evolution of mitochondrial frequencies along the tree:

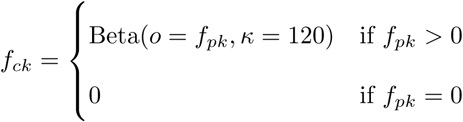

Resampling occurs only when the mutation is present in the parent node (*f_pk_ >* 0).

To simulate read counts given the mutation tree and frequency vectors, we consider *N* cells. For each cell *j*, we randomly select an attachment node *i* from the mutation tree. The mutation frequencies *f_ik_*at node *i* represent the frequencies of each mutation *k* for cell *l*. We simulate read counts 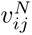 for each nucleotide *N* (Reference *R*, Mutation *X*, and alternative nucleotides *Y* and *Z*). The read counts are drawn from the following multinomial distribution:

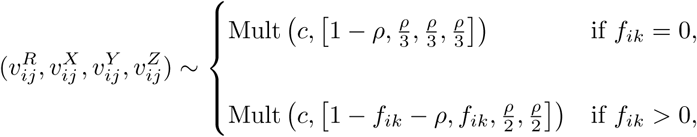

where *c* is the sequencing coverage, and *ρ* the sequencing error rate.

We perform the data simulation for 10 different replicates for all parameter combinations given in Table 1.

**Table 1:** Parameters and values used in simulating the benchmark data.

| Parameter description | Symbol | Parameter sets |
| --- | --- | --- |
| Number of mutations | $L$ | {10, 50} |
| Number of cells | $N$ | {100, 500} |
| Error rate | $\rho$ | {0.05, 0.005, 0.0005} |
| Sequencing coverage | $c$ | {500, 50} |
| Initial mutation frequency | $f_{\text{init}}$ | [1%, 10%] |
| Concentration parameter | $\kappa$ | 120 |

### Inference

To benchmark mt-SCITE against existing approaches, we compare it to MERLIN [34], as well as the standard heuristic pipeline of mutation calling followed by distance-based tree reconstruction using Neighbour-Joining (NJ) [39] and Unweighted Pair Group Method with Arithmetic Mean (UPGMA) [40]. We additionally include SCITE [38] as a baseline.

For MERLIN, we provided a total read count matrix and a variant read count matrix, both with dimensions corresponding to the number of mutations by the number of cells. For our simulated data, the total read count matrix simply contains the sequencing coverage for every mutation and cell, i.e. 500 and 50 (see Tab. 1). The variant count matrix was generated by summing the read counts of the three non-reference nucleotides for each mutation in each cell.

SCITE requires a binary matrix of dimensions corresponding to the number of mutations by the number of cells, where the presence (1) or absence (0) of a mutation in a cell is specified. We defined a mutation as present in a cell, if any of the non-reference nucleotides had read count ≥ 1. Since the true error rate in an experiment is unknown, we ran SCITE with 3 different false positive error rates, {5e-2, 5e-3, 5e-4}, to evaluate its performance under correctly and incorrectly specified error rates. We ran SCITE for 10^6^ iterations and used the first maximum a posteriori (MAP) tree reported for performance comparison.

For NJ and UPGMA, we imitate standard practice by first calling mutations and then performing tree reconstruction. We adopt a simple strategy, as used previously [34], namely we call a mutation as present in the cell if its variant allele frequency exceeds 0.05. Because our simulated per-base error rates span 5e-4 to 5e-2 (Tab. 1), this threshold allows us to evaluate how the standard pipeline degrades as the error rate approaches the calling threshold. From the resulting binary cell-by-mutation matrix, we reconstruct the distance matrix based on pairwise Hamming distances and reconstruct the cell lineage tree using UPGMA (SciPy v1.16.2, scipy.cluster.hierarchy.linkage with method=’average’ [50]), and NJ (Biopython v1.87, Bio.Phylo.TreeConstruction.DistanceTreeConstructor [51]).

For mt-SCITE, we evaluated its performance using the learned error rate. The error rate was learned by performing 3-fold cross validation with 3 repetitions. Specifically, we allow mt-SCITE to select from 20 logarithmically spaced error rates spanning one order of magnitude above and below the true value. Similar to SCITE, mt-SCITE was run for 10^6^ iterations, and the first reported ML tree was used for performance comparison.

### Metrics

We assess two aspects of tree reconstruction quality: the mutation tree topology (for SCITE, mt-SCITE, and MERLIN; UPGMA and NJ do not produce mutation trees) and the cell lineage tree (for all methods). To evaluate the structure of the estimated mutation trees, we compared them to their respective simulated ground truth tree using the parent-child-distance [41].

Cell lineage accuracy was assessed using the root mean squared error (RMSE) between the true and estimated cell-cell distance matrices for each method. Ground truth cell-cell distances are defined as the number of edges on the path connecting each cell pair’s attachment points in the simulated mutation tree. The distances for mt-SCITE and SCITE are computed equivalently on their estimated mutation trees. For MERLIN, we compute the cell lineage tree from the binarised U matrix using its built-in function and compute cell-cell distances as patristic distances on that tree. For NJ and UPGMA, we likewise use patristic distances on the reconstructed cell lineage tree. Note that the scales of the distances are comparable as NJ and UPGMA branch lengths from Hamming distances are effectively mutation counts.

Both metrics are reported as mean ± standard deviation across 10 replicates per error rate.

### Experimental dataset

To evaluate all methods on experimental data with a known ground-truth lineage, we used the TF1 subcloning experiment of Ludwig et al [21]. In this dataset, single cells of the hematopoietic TF1 cell line were isolated, clonally expanded into subclonal populations, and bulk ATAC-sequenced over eight generations, yielding a lineage tree with experimentally defined topology. We analysed 65 subclonal populations together with two bulk samples of the founding population, for 67 samples in total. The original authors called a set of 44 confident variants with their variant-calling pipeline, which we reuse here.

mt-SCITE, SCITE, and MERLIN all make the infinite-sites assumption (ISA), under which a variant that is already present in the founding population can drift below and above a variant calling threshold and leave spurious signal for a lost or regained mutation. We therefore remove such variants. For MERLIN and SCITE, which operate on binarised genotypes rather than per-cell mutation probabilities, we filter out ancestral variants by their variant allele frequency (VAF) in either bulk sample, and apply two thresholds as a sensitivity analysis: a 5% threshold, which removes 2 variants (7790 C and 8003 T), leaving 42 variants; and a 1% threshold, which removes 6 variants (7790 C, 8003 T, 4038 A, 8207 A, 309 T, and 13709 A), leaving 38. For each method we report results both on the full 44-variant set and on the filtered sets. For mt-SCITE we additionally reuse the filtering pipeline on the level of the mutation probability matrix from our other experimental analyses, which removes variants whose probability of being mutated in the bulk samples exceeds 0.9 (*P* (mut) *>* 0.9).

Neighbour-joining (NJ) and UPGMA do not assume infinite sites, so ancestral variants do not violate their models; we therefore retain the full variant set for these methods. We binarise the variant calls at VAF 5%, compute pairwise Hamming distances between cells, and apply NJ and UPGMA to the resulting distance matrices.

## Data analysis

### Data availability

In this manuscript, we analysed four publicly available datasets. The first dataset, based on ATAC-sequencing data, was generated from 6 *in vitro* expanded CD8+ T-cells with two replicates per clonal expansion resulting in a total of 12 samples. Note that in each sample there were 500 - 1000 cells which were bulk sequenced. The ATAC-seq dataset is available from [36].

The second dataset is based on scRNA-seq data derived from T-cells from a healthy human donor vaccinated against Yellow Fever Virus. It consists of 72 Smart-seq2 libraries generated from CD8+ T-cells and is available from [35, 37]. We further use a bulk sample comprising PBMCs collected from the same individual which were subjected to WGS to identify and filter out germline mutations. The WGS data is available from [52].

Third, we analysed a large scale single cell mitochondrial genotyping dataset from MAESTER [22] of a bone marrow aspirate from a human donor with clonal hematopoiesis, containing 9346 cells. The dataset analysed in this manuscript consists of mitochondrial variant calls obtained by mitochondrial variant enrichment from high-throughput single-cell RNA sequencing of sample BPDCN712, comprising 1,397 cells with at least 1% VAF for one of 26 informative mitochondrial variants. The MAESTER data is available from the Gene Expression Omnibus under accession GSM5534706.

Fourth, we analysed a dataset with an experimentally defined ground-truth lineage: the TF1 subcloning experiment of Ludwig et al. [21]. There, single TF1 cells were isolated, clonally expanded into subpopulations over successive generations, and each subpopulation was bulk ATAC-sequenced, with the known subcloning pedigree serving as the ground truth. We used the 44 mitochondrial variants called by the original study and 67 of the 69 samples, comprising 65 clonally expanded subclones and 2 bulk samples of the founding population (the two “mix” controls were excluded). The deposited data (GSE115208, a subseries of GSE115218; https://www.ncbi.nlm.nih.gov/geo/query/acc.cgi?acc=GSE115208) provide the called variants and their allele frequencies but not per-base read counts required for our analyses. We obtained these per-base read counts directly from Caleb Lareau.

### Sequencing data analysis

Read pairs were trimmed from Illumina adapters using trimgalore. For ATAC-seq data, read pairs were aligned to the human genome (hg38) using STAR and genes annotated using gencode.v26.annotation.gtf.

For scRNA-seq data, read pairs were aligned to the human genome (hg38) using BWA-mem [53]. BAM files were sorted and indexed using samtools. Read counts supporting each nucleotide at each position in the mitochondrial genome were obtained by allelecounter (https://github.com/cancerit/alleleCount). In scRNA-seq data, clonotype analysis was performed by mixcr [54]. We included TCR clones that showed support for at least one mitochondrial mutation based on VAF. The procedure for cell type classifications of the analysed CD8+ T-cells subjected to scRNA-seq is described in [35].

We selected clones (defined by TCR) that showed support for at least one mitochondrial mutation (based on VAF). This was done deliberately to evaluate the method on real data with known ground truth since we needed a dataset that actually contained a mitochondrial mutation signal. We agree that, ideally, one would select cells without conditioning on mutation status. However, in the absence of any mutation signal, tree inference is not possible, and the method’s performance cannot be assessed. Selecting clones with at least one supported mutation therefore represented a reasonable trade-off. This design still allowed us to (a) test whether the algorithm could re-identify the known clones and (b) identify additional mutations supporting the splits, as well as subclonal mutations.

For the MAESTER dataset, variant selection followed the filtering criteria of the original study: variants were required to have mean coverage above 5, mean base quality of at least 30, to be absent in more than 90% of cells (likely germline mutations), and to be present at greater than 50% variant allele frequency in at least 10 cells.

### Phylogenetic inference using mt-SCITE on experimental data

To compute mutation probabilities for the ATAC-seq and scRNA-seq datasets, we selected a range of error rates based on prior studies that did not use mitochondrial mutation data, essentially using these rates as a plausible prior distribution. Specifically, we used error rates ranging from 0.0001 to 0.0351 for the ATAC-seq data [35], and from 0.01 to 0.23 for the scRNA-seq dataset [36]. For the MAESTER dataset, we used error rates ranging from 1e-5 to 0.1, based on the minimum mean base quality of 30 which corresponds to an a priori error rate of 10^−3^ and allowing for at least one order of magnitude variation in each direction. For each error rate, we generate a mutation probability matrix using mt-SCITE (Methods) and then applied several filtering steps to remove mutations unlikely to be part of the mutation tree.

First, we remove sites that are likely germline mutations. For the first two datasets, we achieve that by filtering variants with a mutation probability exceeding 90% in an unamplified bulk sample from a mixture of cells from the same donor. For the MAESTER dataset, germline variants are already filtered out before mutation probability matrix calculation. Second, we discard sites that do not have a mutation probability greater than 90% in at least 2 cells. Third, we remove sites where 50% or more of the cells have missing data, where a site is defined as missing in a given cell when its total read coverage at that position is zero (*c_ij_* = 0), or the reference nucleotide at that position is undefined (encoded as N). For the remaining sites, the missing values are set to the prior mutation probability. Fourth, we only retain those sites that do not have another mutated site within the next 4bp to reduce false positive mutations that may stem from alignment artifacts or local sequence complexity.

For all datasets, we apply the error learning procedure using 3-fold cross validation with 3 repetitions and report the learned error rate. For the first two datasets, we require trees to have at least 7 mutations, given that we have data from 6 clones and 1 bulk sample and expect to observe at least one defining mutation per clone.

Independently, we run mt-SCITE with 10 different random seeds to start the MCMC procedure from different starting states. For every filtered mutation probability matrix, we keep the list of co-optimal maximum likelihood trees across these runs. Unlike the trees obtained through the error learning procedure, these trees are estimated from the full data set.

On this set of trees, we compute the average tree purity and clone coverage (Methods) for every error rate.

## Author Contributions

Conceptualization: J.H., Methodology: J.H., P.F.F., S.S., J.K., K.J., P.B., Software: J.H., S.S., P.F.F., K.J., P.B., C.J.E., Validation: J.H., S.S., Formal Analysis: J.H., S.S., P.F.F., Investigation: J.E.M., J.M., Data Curation: J.H., S.S., Writing-Original: J.H., S.S., Writing-Review & Editing: all authors, Visualization: J.H. S.S., Supervision: J.K., N.B., Project Administration: J.H., S.S., Funding acquisition: N.B. S.S. and J.H. contributed equally to this work as co-first authors. When referencing this paper in their CVs or other professional materials, each co-first author may list their own name first.

## Acknowledgments

The authors thank Gordon J. Köhn for advice on the Snakemake workflow and Caleb Lareau for generously sharing the per-base read count data for the TF1 subcloning experiment.

