## supplemental-materials for "Probabilistic cell lineage inference from mitochondrial mutations"

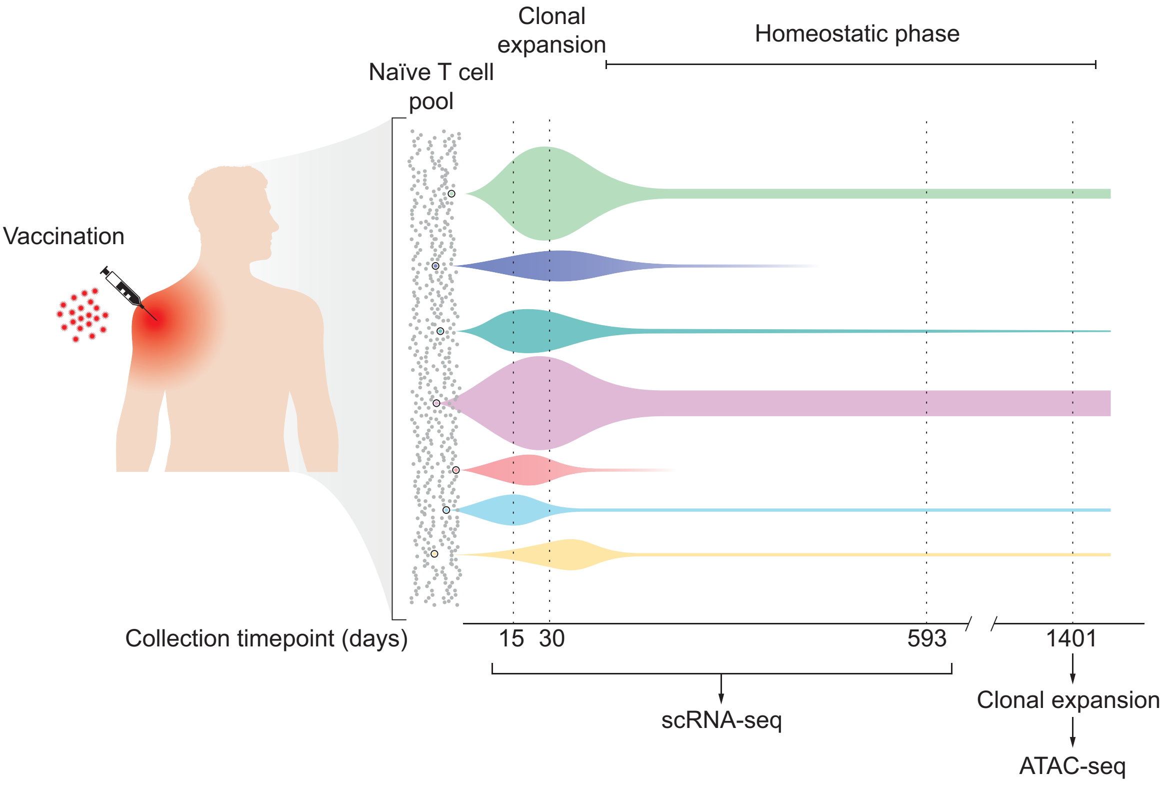


**Fig. 1**: Experimental setup. Human donors were vaccinated against YFV. The vaccination triggers the activation of naïve CD8+ T cells which undergo a rapid clonal expansion during the acute phase of the immune response followed by a slower homeostatic division. Single CD8+ T cells are isolated from blood at different timepoints after vaccination. Cells isolated at day 1,401 post-vaccination were expanded in vitro and subjected to ATAC-seq. Cells isolated at day 15, 30 and 593 were subjected to scRNA-seq.

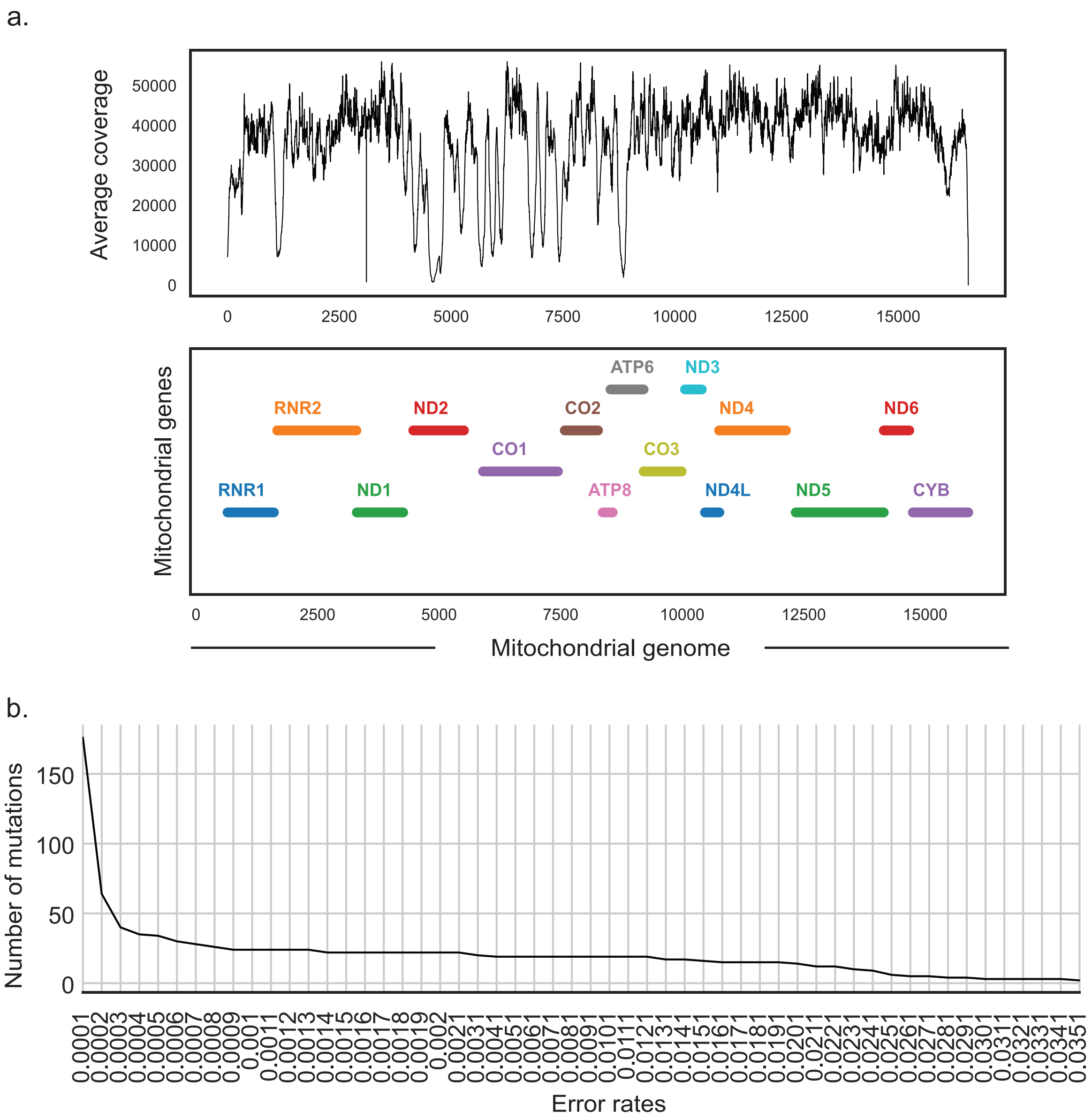


**Fig. 2**: Tree inference of clonal expansions subjected to ATAC-seq (a) The average sequencing depth over mitochondrial genome positions and genes. (b) The number of detected mutations.

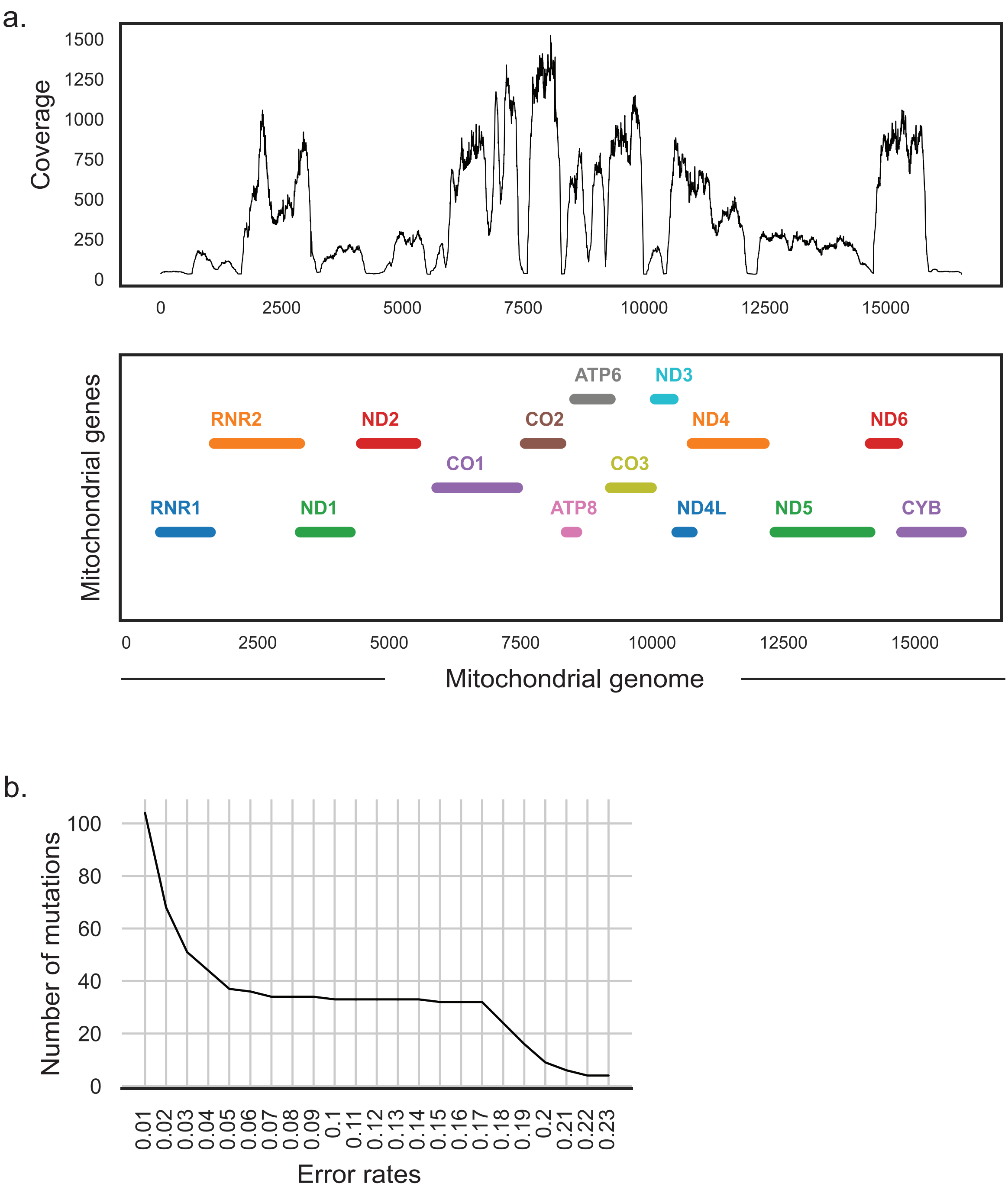


**Fig. 3**: Tree inference of in vivo expanded CD8+ T cells subjected to scRNAseq (a) The average sequencing depth over mitochondrial genome positions and genes. (b) The number of detected mutations.

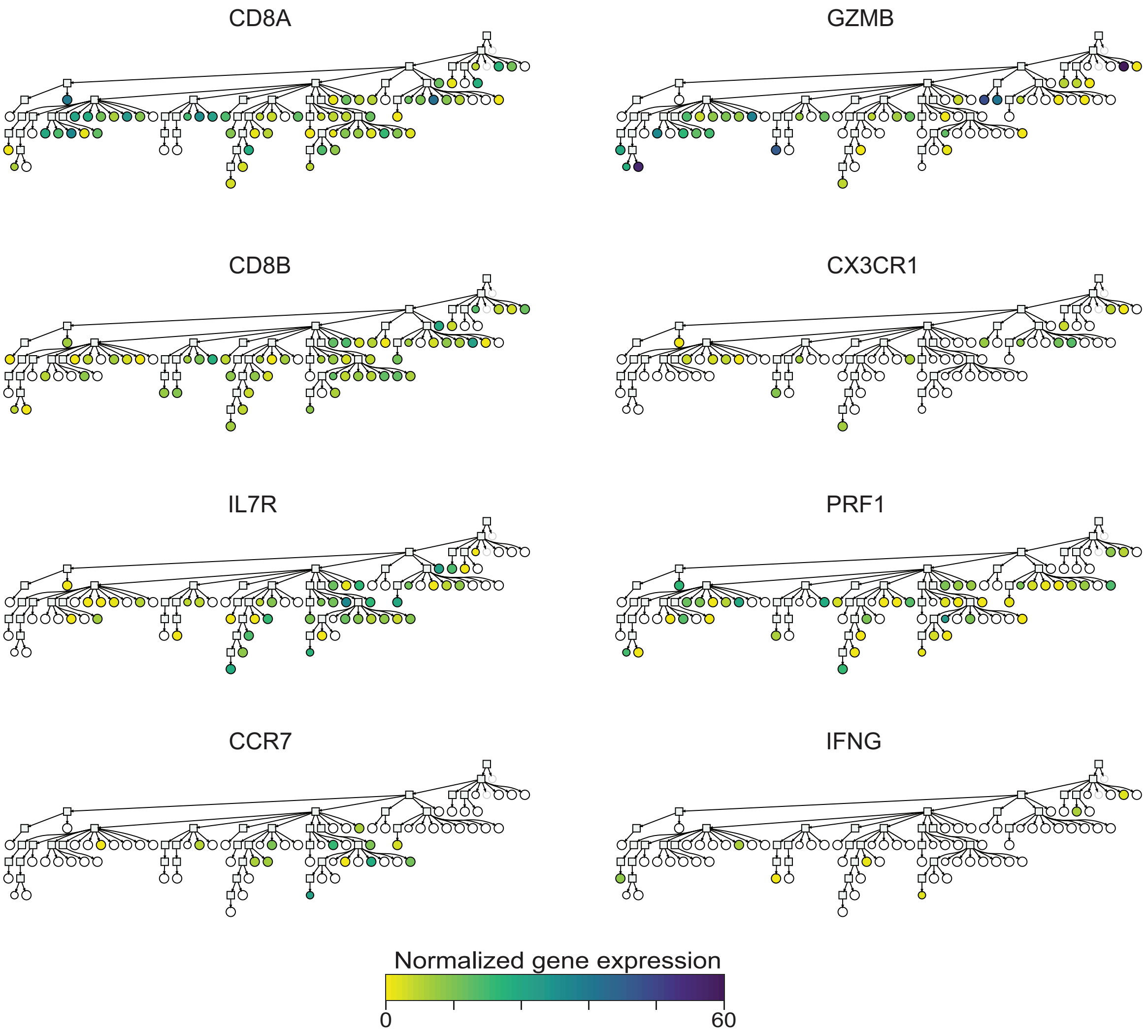


**Fig. 4**: Visualization of gene expression in CD8+ T cells across the phylogenetic tree.

### Additional results for the simulated benchmark

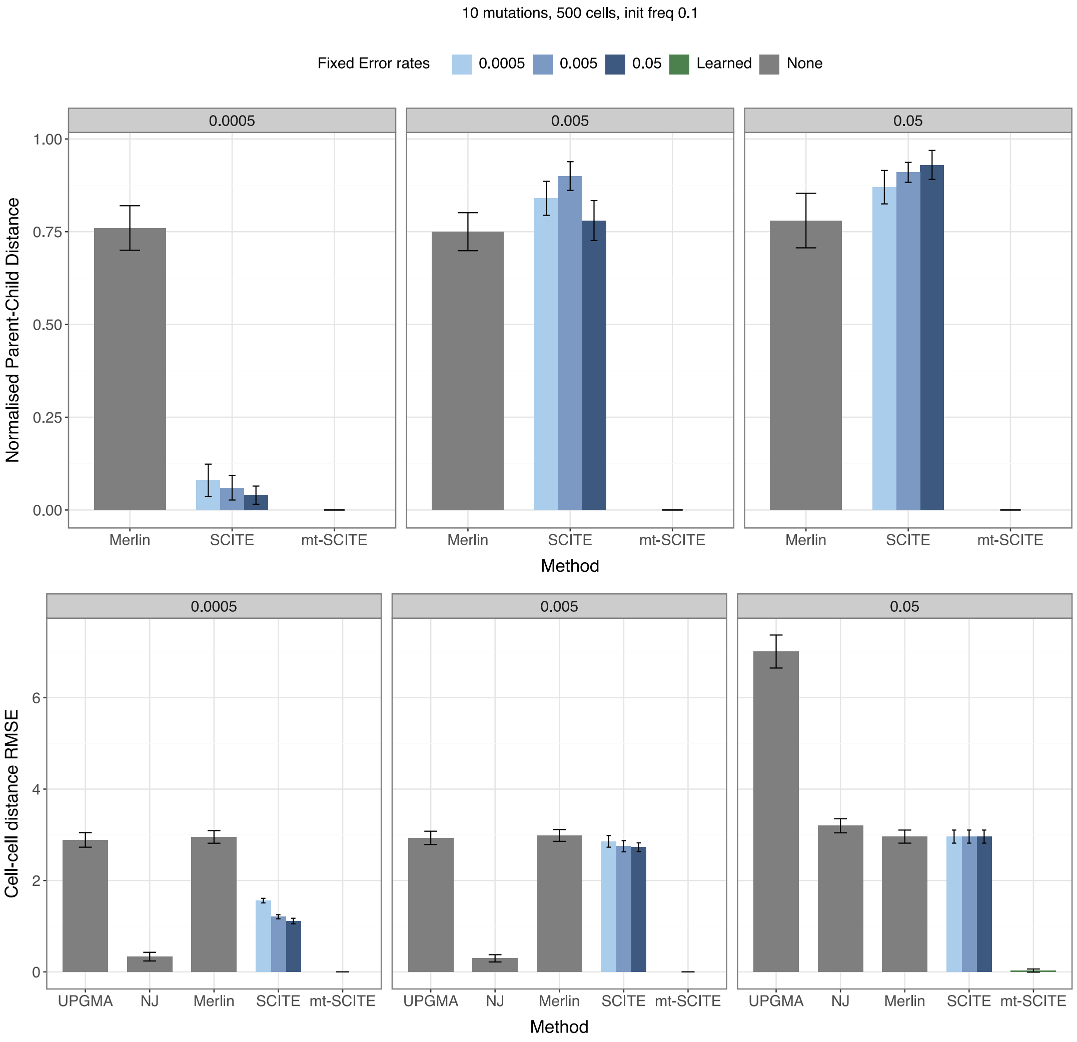


**Fig. 5**: mt-SCITE outperforms existing methods in terms of mutation tree and cell relationship reconstruction on simulated data. Each panel represents input data generated with a true error rate (0.0005, 0.005, and 0.05, left to right). Bars show the normalized parent-child distance (top) and the mean squared error (MSE) between the true and estimated cell-cell distances (bottom), where lower values represent better inference accuracy. Colors denote the error rates used during inference: MERLIN, NJ and UPGMA do not account for error rate ('None'), SCITE was evaluated for three fixed error rates (0.0005, 0.005, and 0.05), and mtSCITE was evaluated for the learned error rate. Results are shown for 10 mutations, an initial mutation frequency of 10%, and 500 cells.

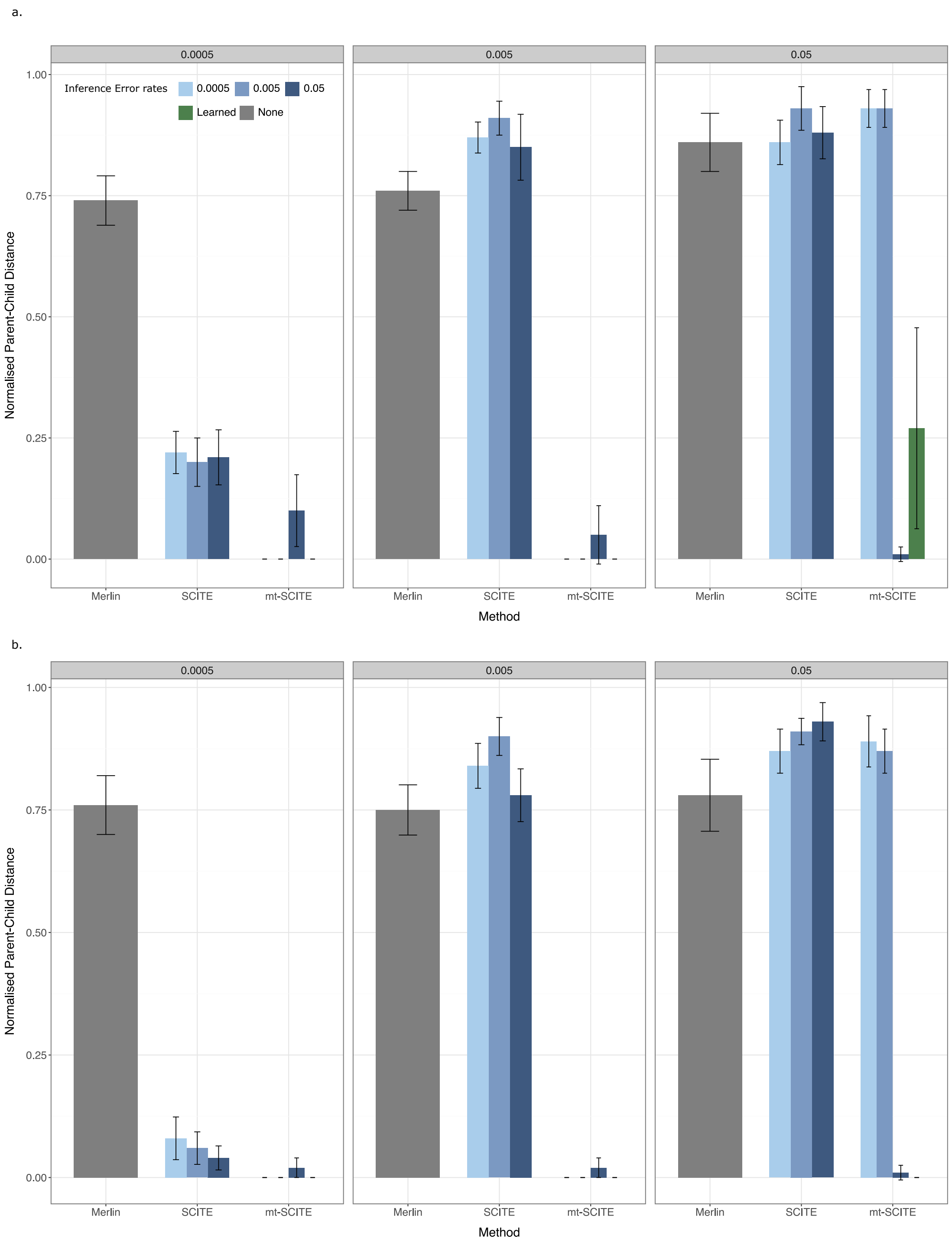


**Fig. 6**: Comparing mt-SCITE for its learned versus fixed error rates. Each panel represents input data generated with a true error rate (0.0005, 0.005, and 0.05, left to right). Bars show the normalized parent-child distance, where lower values represent better inference accuracy. Colors denote the error rates used during inference: MERLIN does not account for error rate ('None'), SCITE was evaluated for three fixed error rates (0.0005, 0.005, and 0.05), and mtSCITE was evaluated for the same fixed error rates and learned error rate. Results are shown for 10 mutations, an initial mutation frequency of 10%, and (a) 100 cells and (b) 500 cells.

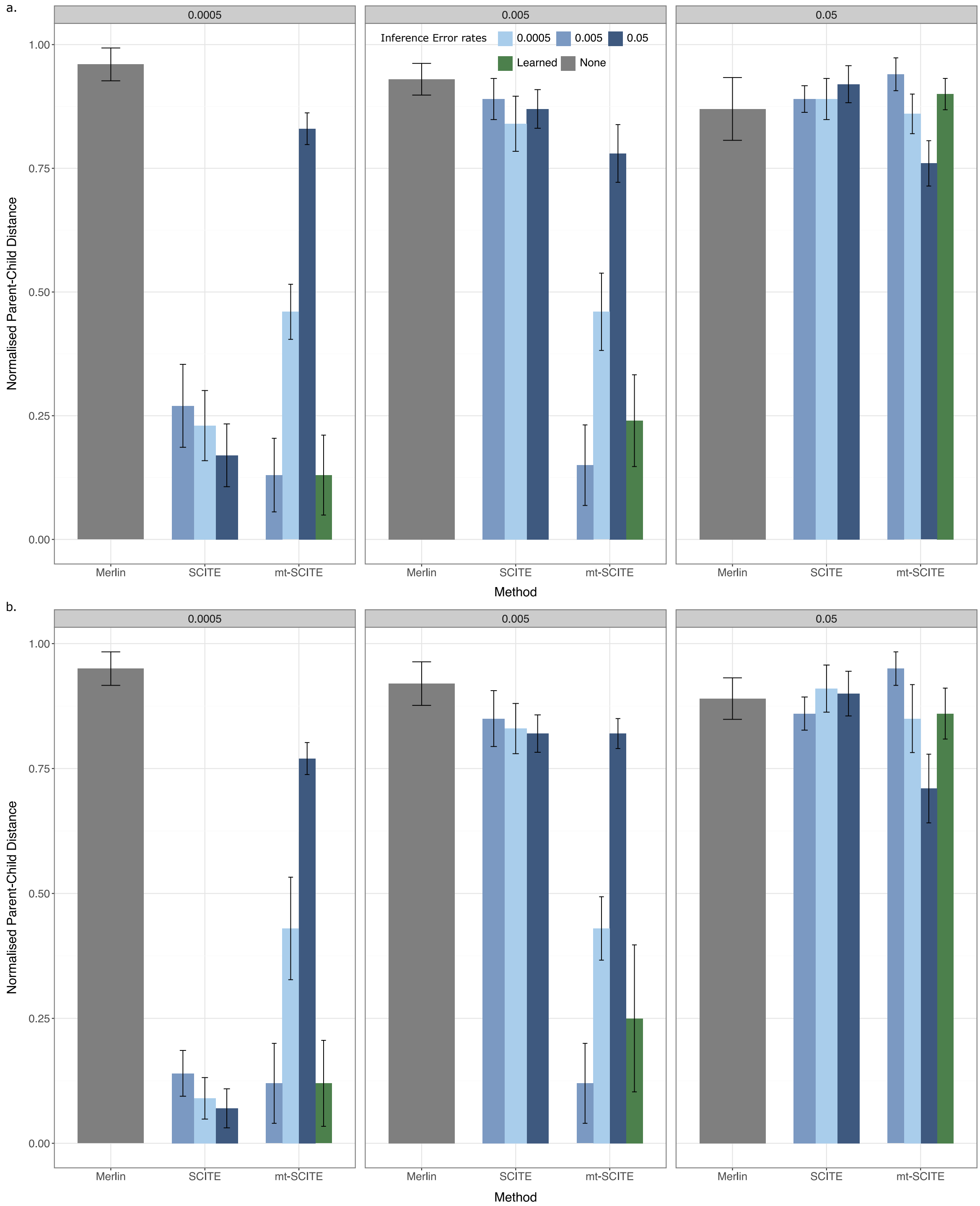


**Fig. 7**: Parent-child distances are shown for 10 mutations, an initial mutation frequency of 1%, and (a) 100 cells and (b) 500 cells. For full caption refer to Supplemental Figure 6.

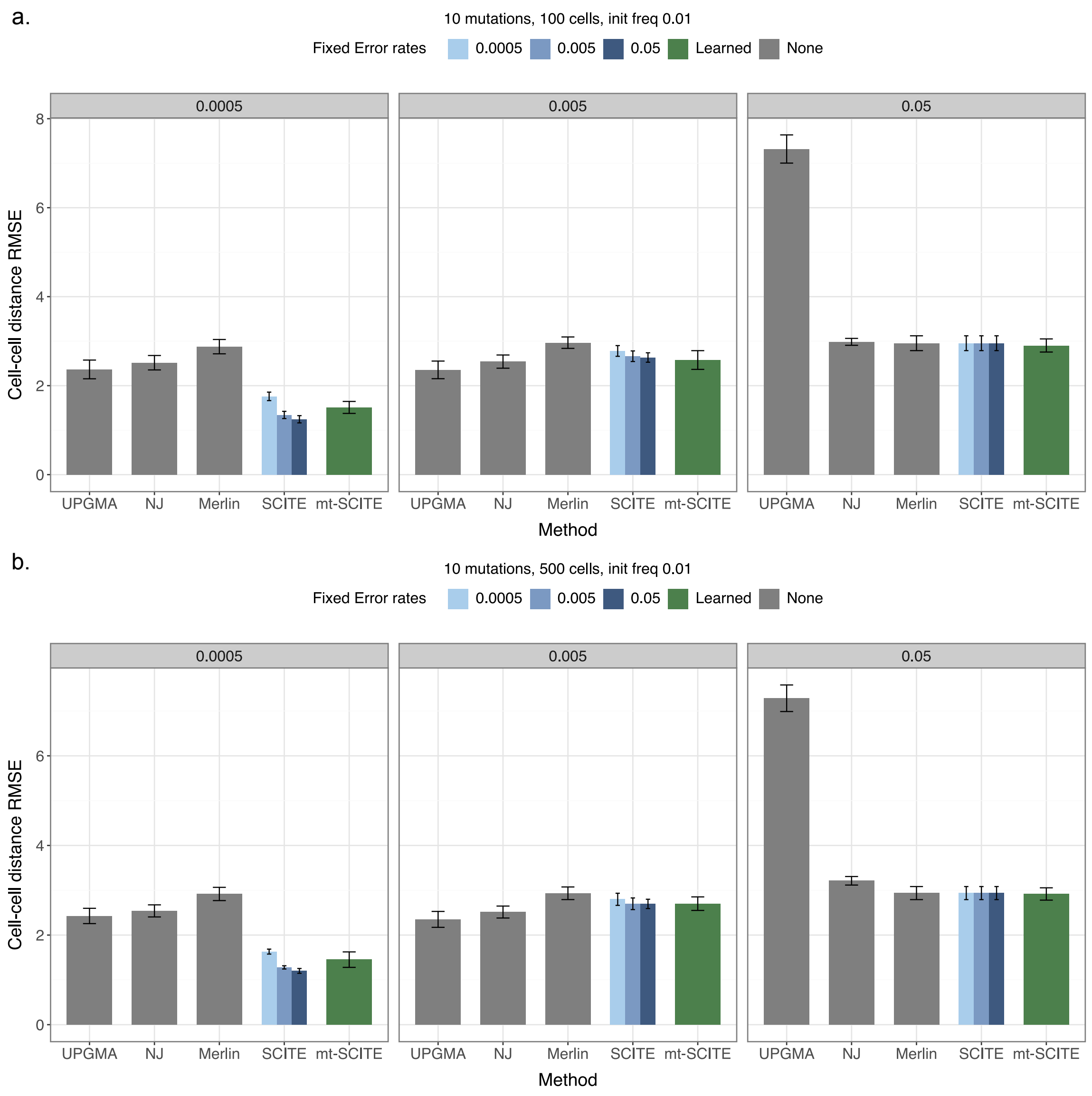


**Fig. 8**: Cell-cell distance errors are shown for 10 mutations, an initial mutation frequency of 1%, and (a) 100 cells and (b) 500 cells.

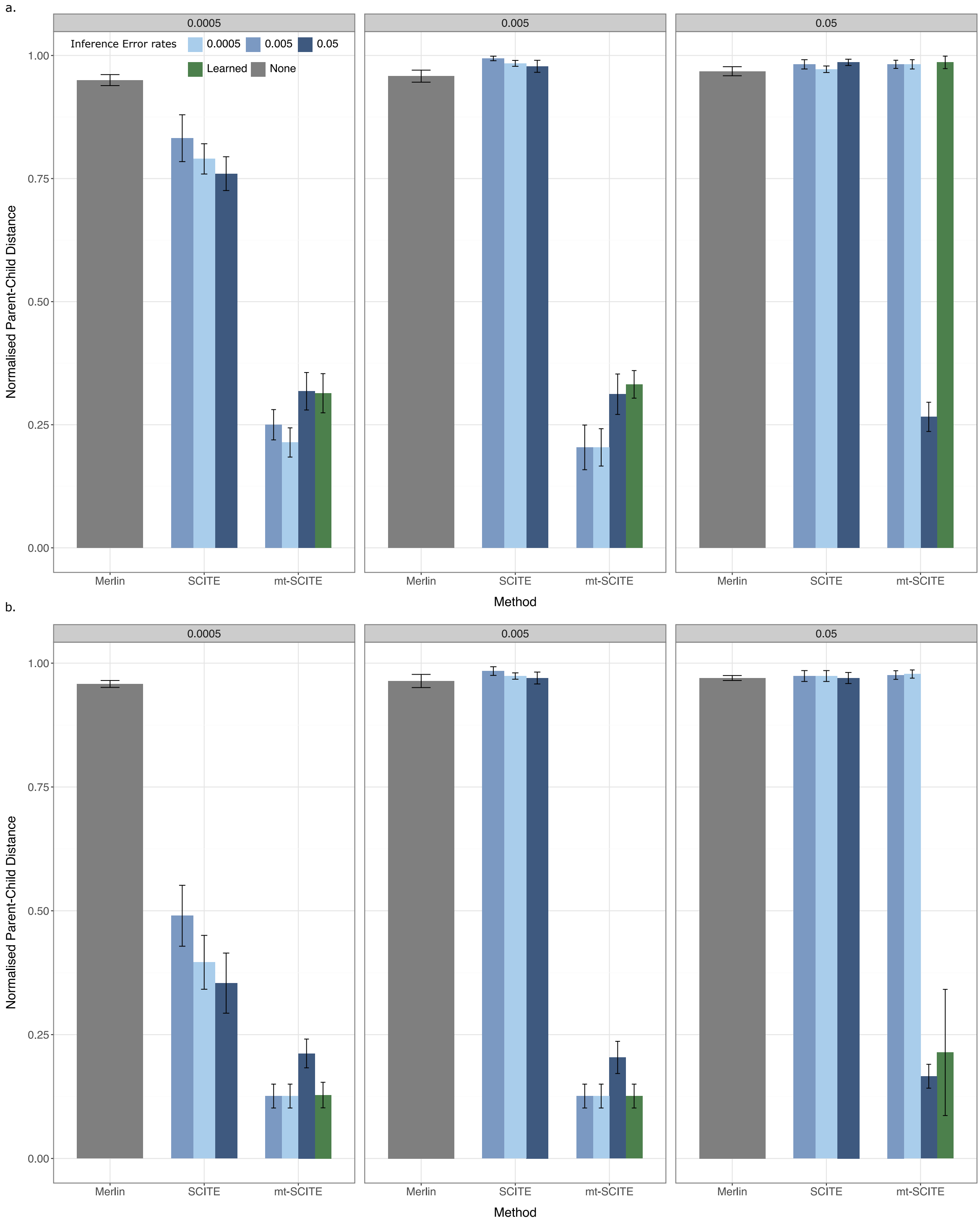


**Fig. 9**: Parent-child distances are shown for 50 mutations, an initial mutation frequency of 10%, and (a) 100 cells and (b) 500 cells.

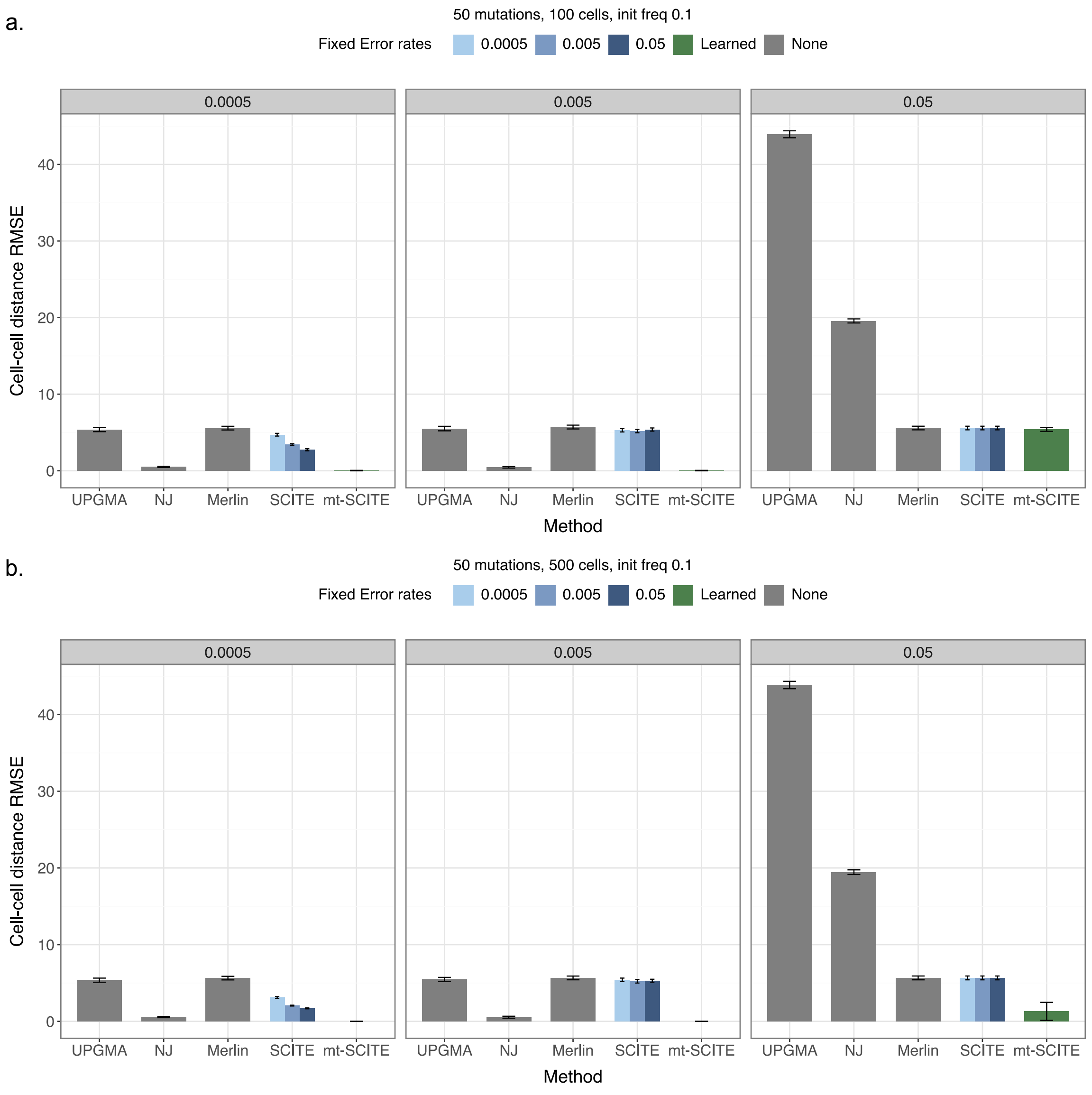


**Fig. 10**: Cell-cell distance errors are shown for 50 mutations, an initial mutation frequency of 10%, and (a) 100 cells and (b) 500 cells.

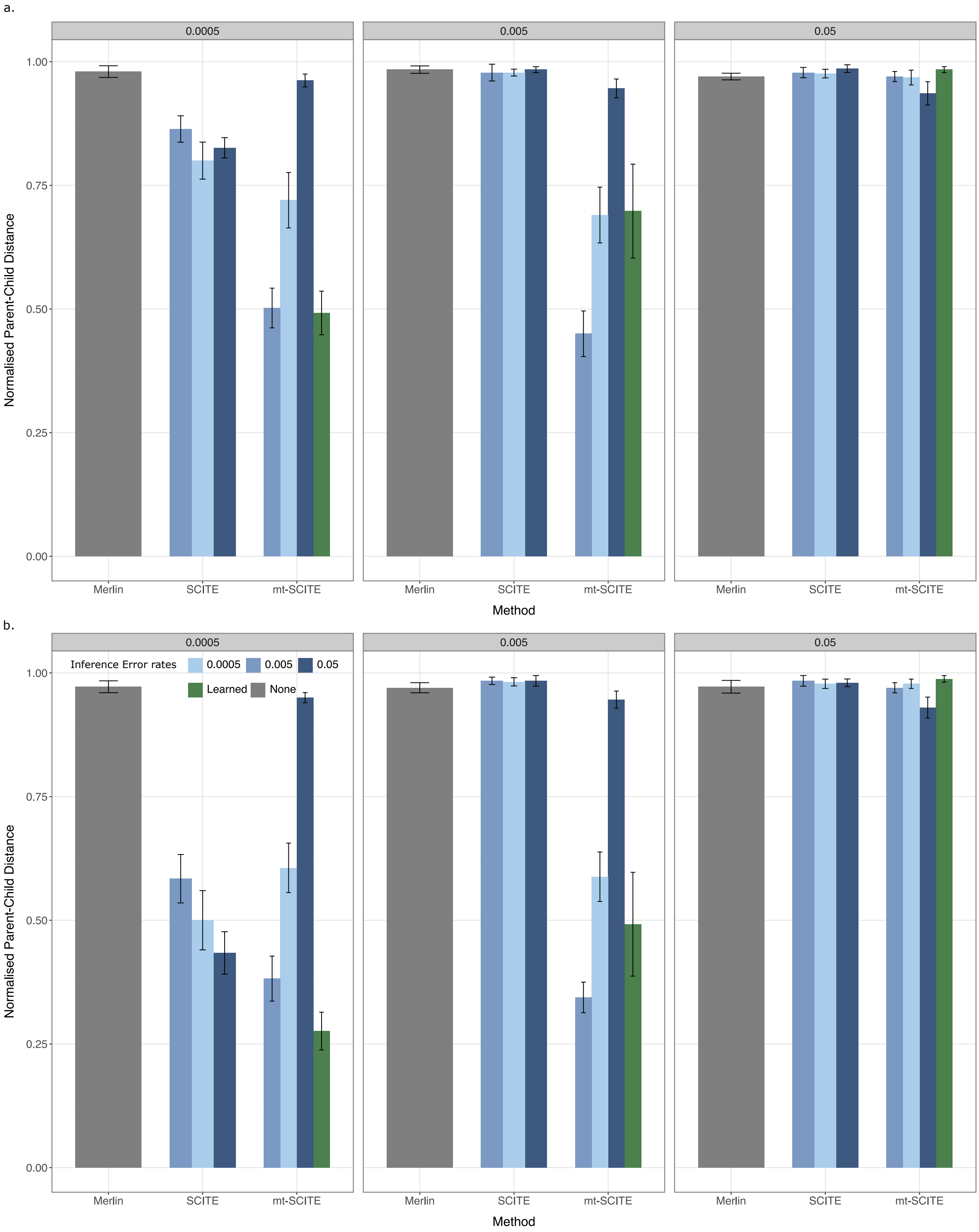


**Fig. 11**: Parent-child distances are shown for 50 mutations, an initial mutation frequency of 1%, and (a) 100 cells and (b) 500 cells. For full caption refer to Supplemental Figure 6.

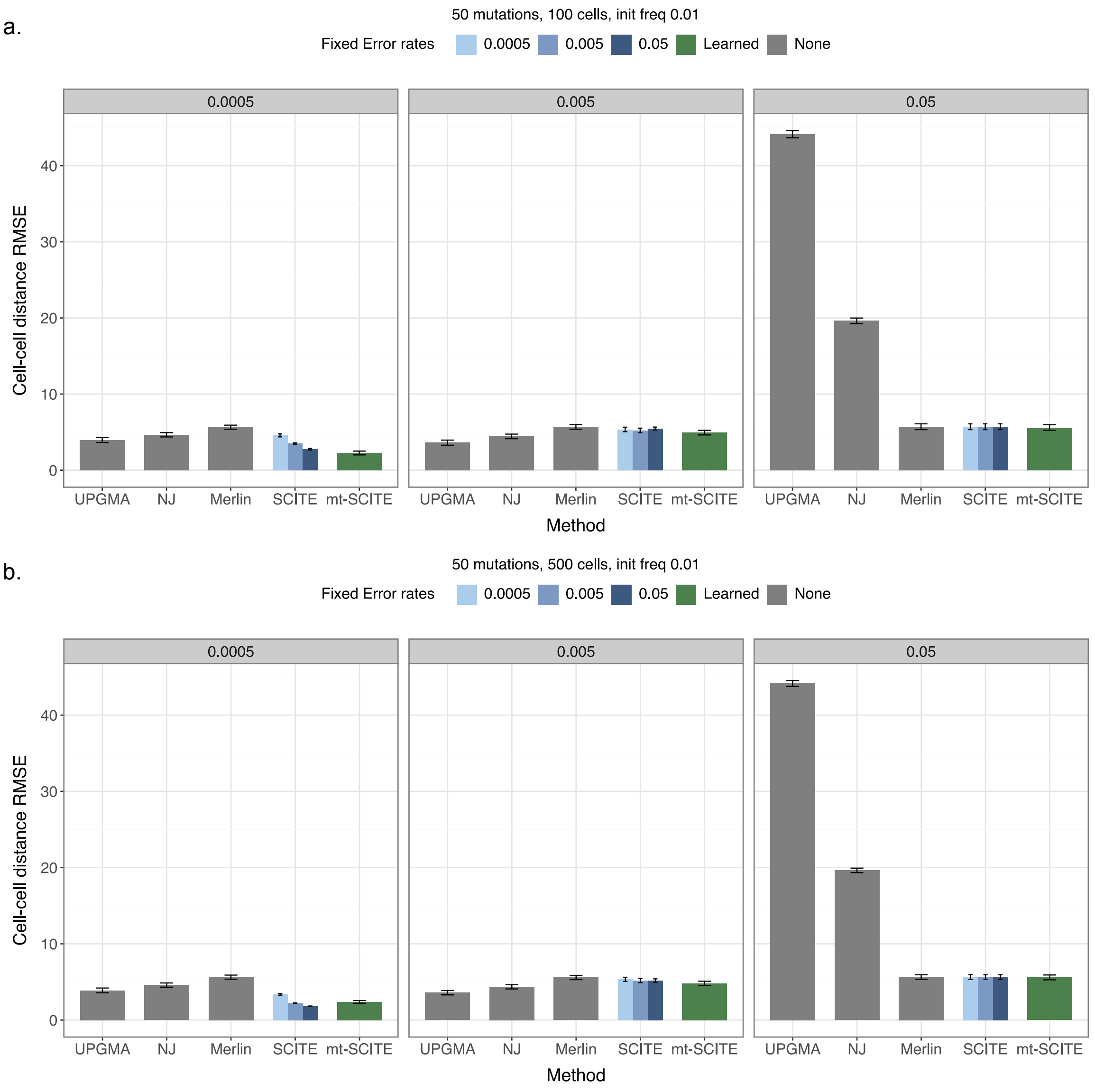


**Fig. 12**: Cell-cell distance errors are shown for 50 mutations, an initial mutation frequency of 1%, and (a) 100 cells and (b) 500 cells. For full caption refer to Supplemental Figure 6.

### Additional results for the benchmark on experimental data

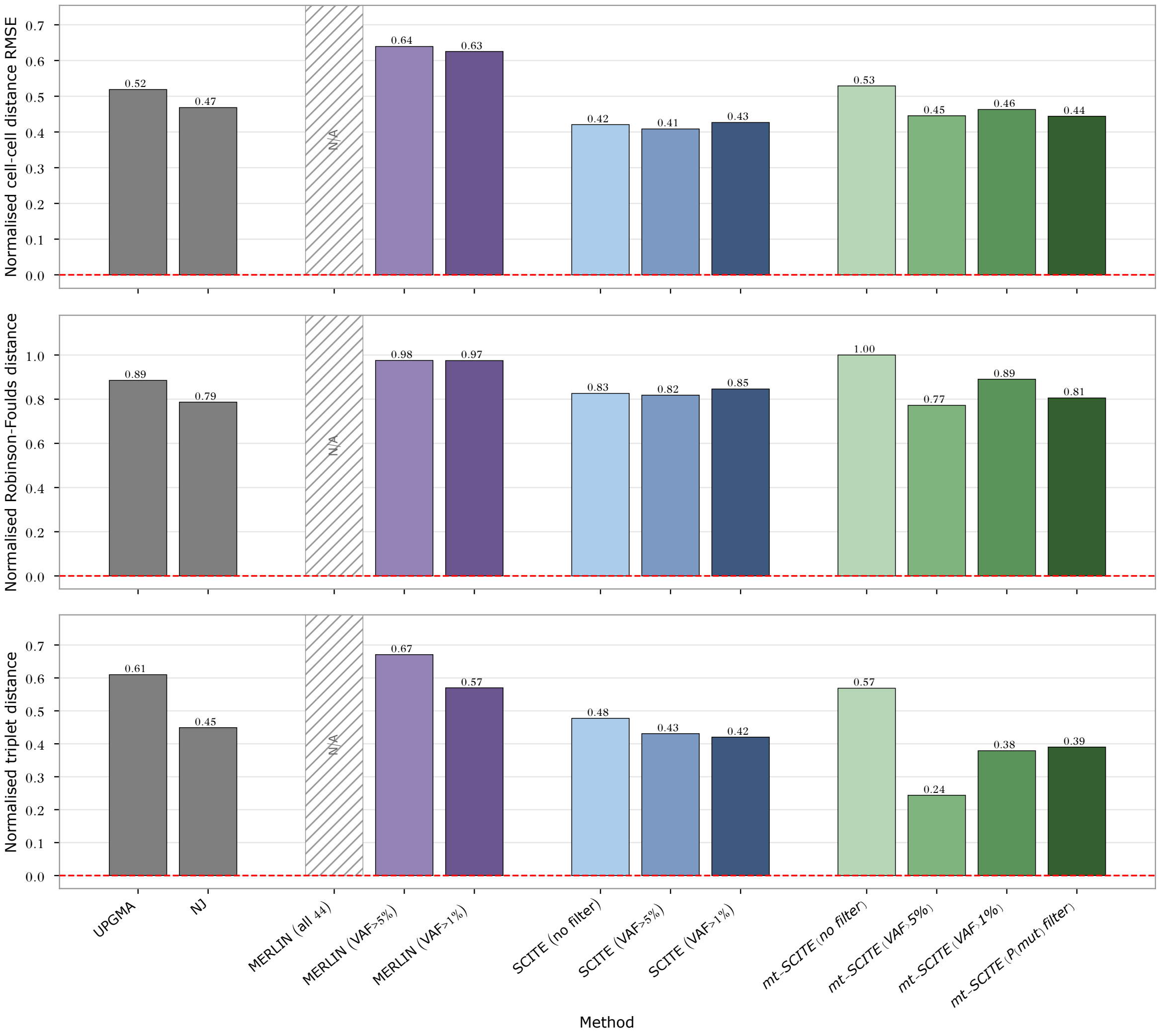


**Fig. 13**: Cell lineage tree topology errors on the TF1 experimental dataset across methods. From top to bottom, we show the normalised cell-cell distance RMSE, Robinson-Foulds distance (RF) and triplet distance for all methods. Methods were run across variant-filtering conditions: no filter, and excluding ancestral variants detected in the bulk samples above 1% and 5% VAF. mt-SCITE was additionally run using its native variant-filtering pipeline based on the mutation probability matrix [P(mut) filter]. Dashed red line indicates zero errors.

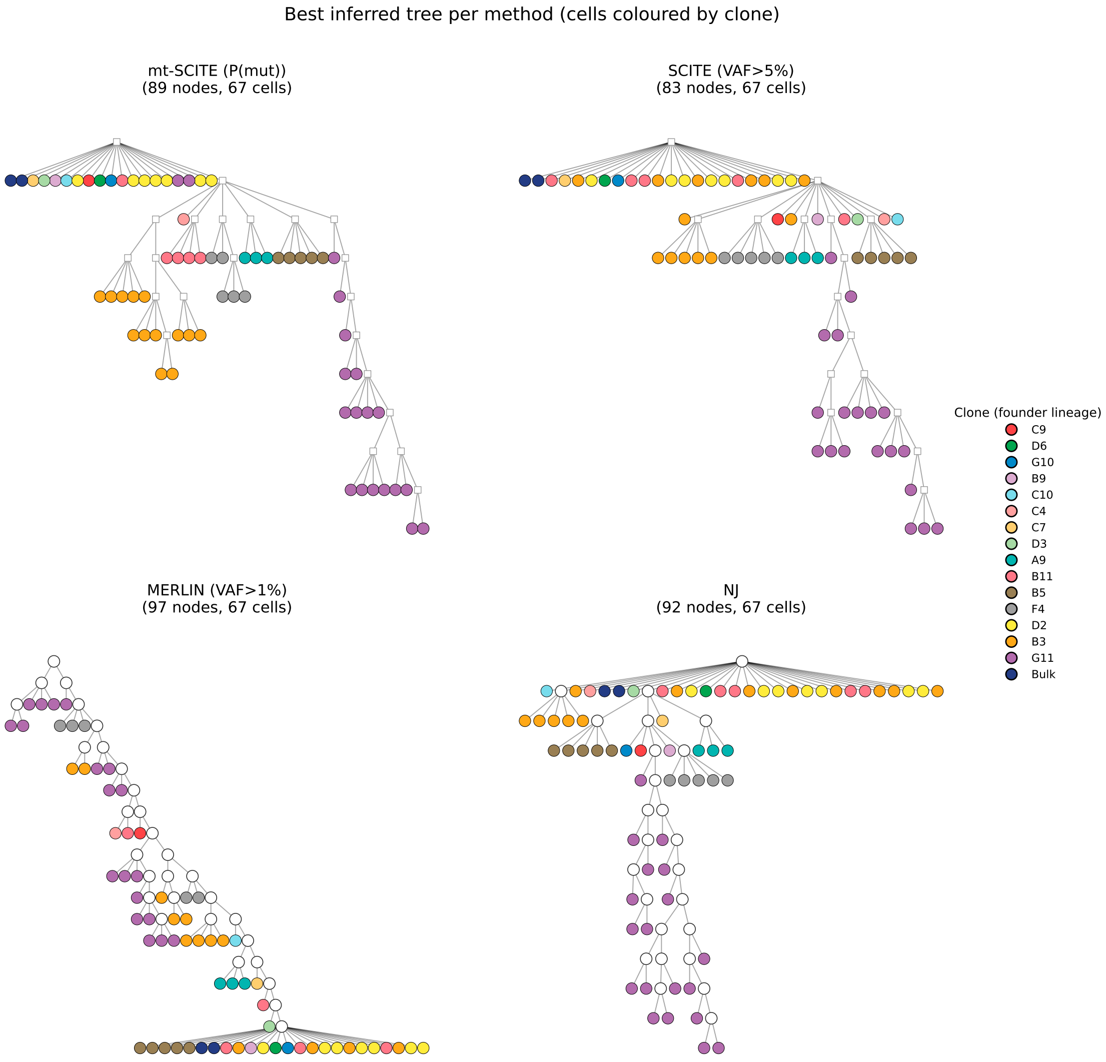


**Fig. 14**: Plotting the best tree per method family on the TF1 experimental dataset.

### Sensitivity analyses

We perform additional sensitivity analyses to show mt-SCITE's behaviour when (a) resampling the heteroplasmy level for every cell (b) the sequencing coverage is lower (50 reads) and (c) when a recurrent mutation occurs. We will show the results for each of these cases in the following subsections.

#### Resampling heteroplasmy levels per cell

We adjust the simulation so that, after drawing each cell's attachment node from the mutation tree, we resample that cell's per-variant heteroplasmy frequencies from the same Beta distributions used along the tree, now centered on the frequencies of its attachment node, before drawing read counts. This adds a further layer of per-cell variation in heteroplasmy relative to the main simulations, modelling cells that share a mutation but are more distantly related. We apply all methods as before and report results in Figures 15 - 18.

We find that mt-SCITE is robust at low-to-moderate error rates and even benefits from increased variance in per-cell heteroplasmy. Relative to the main simulations, its accuracy is maintained or substantially improved across initial frequencies (1% and 10%) and cell numbers (100 and 500) at the lower error rates (0.0005 and 0.005); the gains are largest at the low 1% initial frequency, where the normalised parent–child distance falls essentially to zero and the cell–cell distance RMSE decreases by ∼67–74%, while at the 10% initial frequency, where reconstruction is already near-perfect under both models, accuracy is essentially unchanged. Across this low-to-moderate error regime mt-SCITE is the single most accurate method in every corner for both cell–cell distance and mutation-tree reconstruction. At the highest error rate (0.05) the added per-cell variance decreases performance: mt-SCITE's accuracy degrades relative to the main simulations and shows greater variance. It nonetheless remains the most accurate method for cell–cell distance reconstruction at every setting, and for mutation-tree reconstruction at the 10% initial frequency; only for mutation-tree reconstruction at the low 1% initial frequency does MERLIN marginally surpass it (by ∼5–11% in normalised parent–child distance), while mt-SCITE retains its cell–cell distance advantage there. In this high-error regime the benefit of additional cells is clearest at the 10% initial frequency, where increasing from 100 to 500 cells substantially reduces the cell–cell distance RMSE. This pattern is expected: because mt-SCITE pools information across all cells attached to the same mutation node, greater spread in per-cell heteroplasmy means some cells sample the variant at higher frequency, and these more confident observations are combined to support the shared mutation. This advantage that is strongest when the underlying signal is weak (low initial frequency) and that only the highest error rate overwhelms.

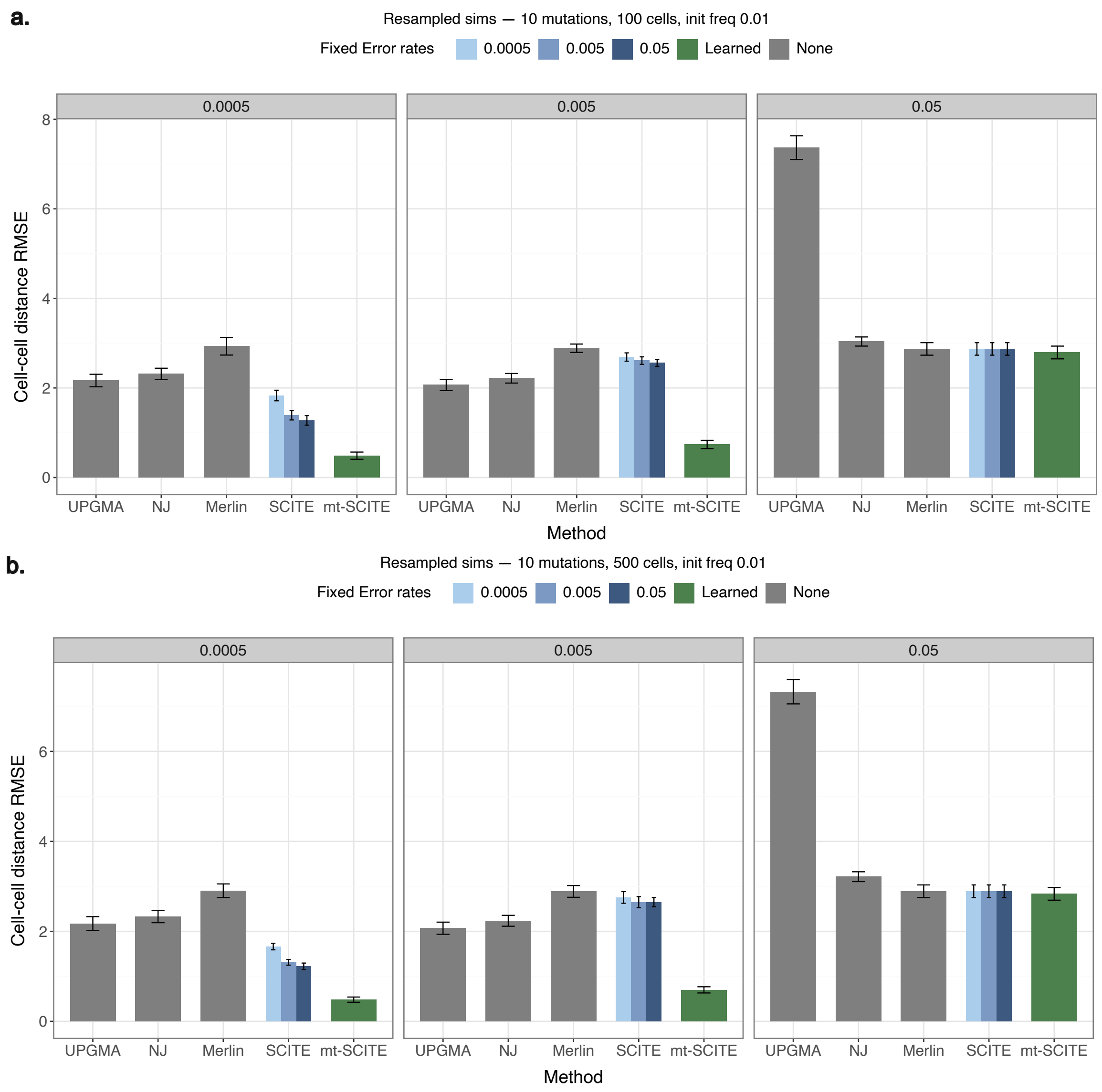


**Fig. 15**: Cell-cell distance errors for simulations with cell heteroplasmy level resampling are shown for 10 mutations, an initial mutation frequency of 1%, and (a) 100 cells and (b) 500 cells.

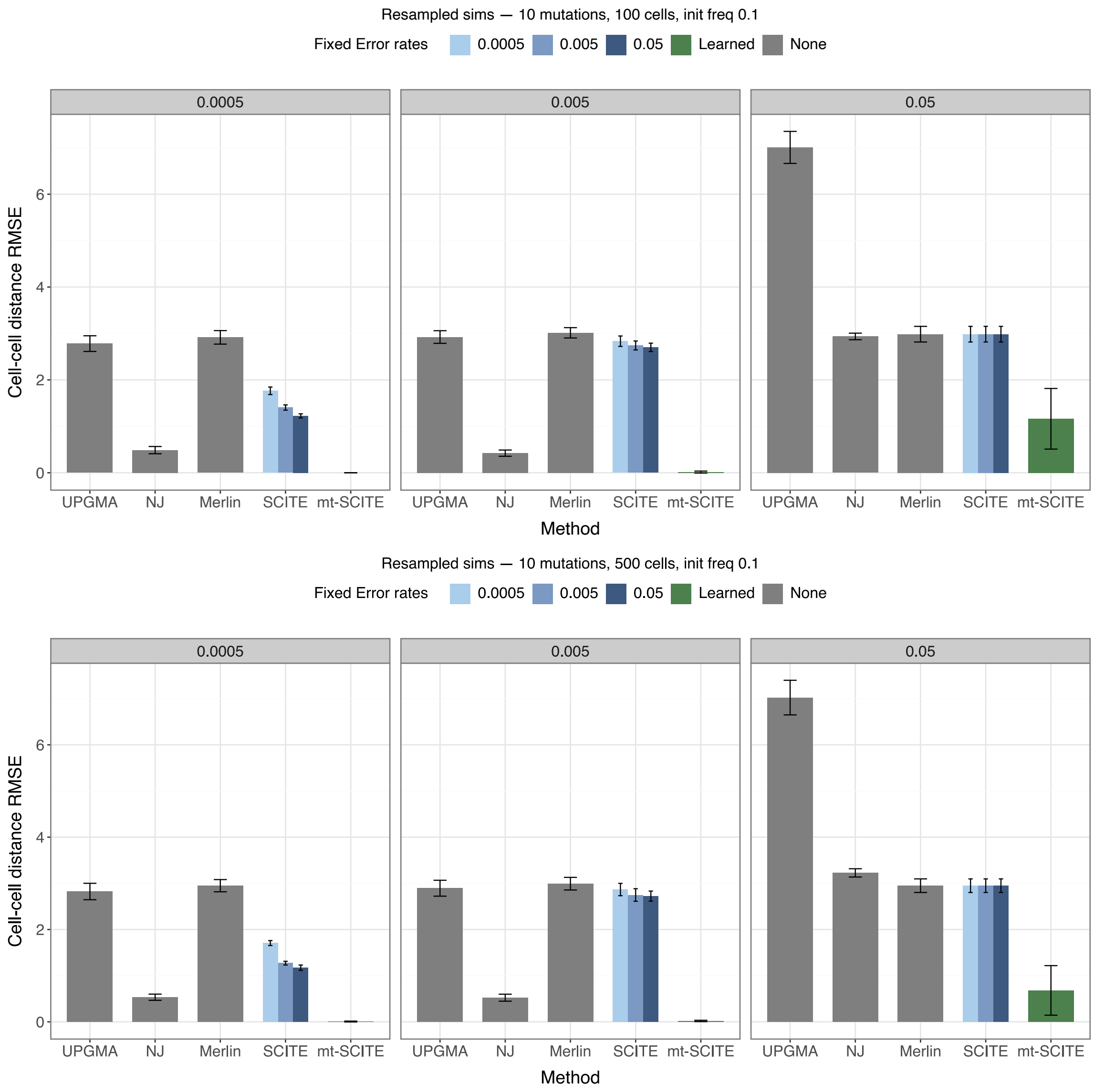


**Fig. 16**: Cell-cell distance errors for simulations with cell heteroplasmy level resampling are shown for 10 mutations, an initial mutation frequency of 10%, and (a) 100 cells and (b) 500 cells.

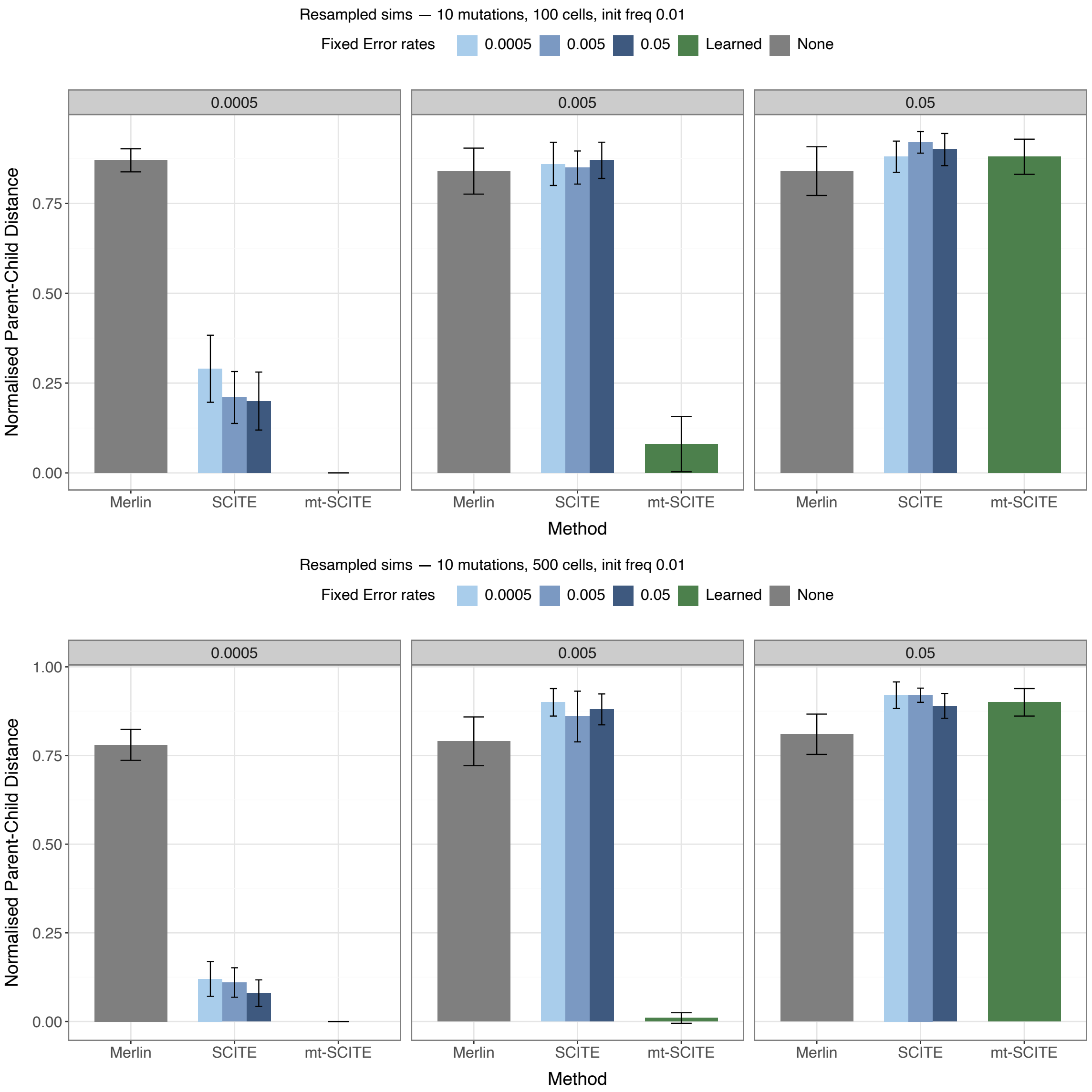


**Fig. 17**: Parent child distances for simulations with cell heteroplasmy level resampling are shown for 10 mutations, an initial mutation frequency of 1%, and (a) 100 cells and (b) 500 cells.

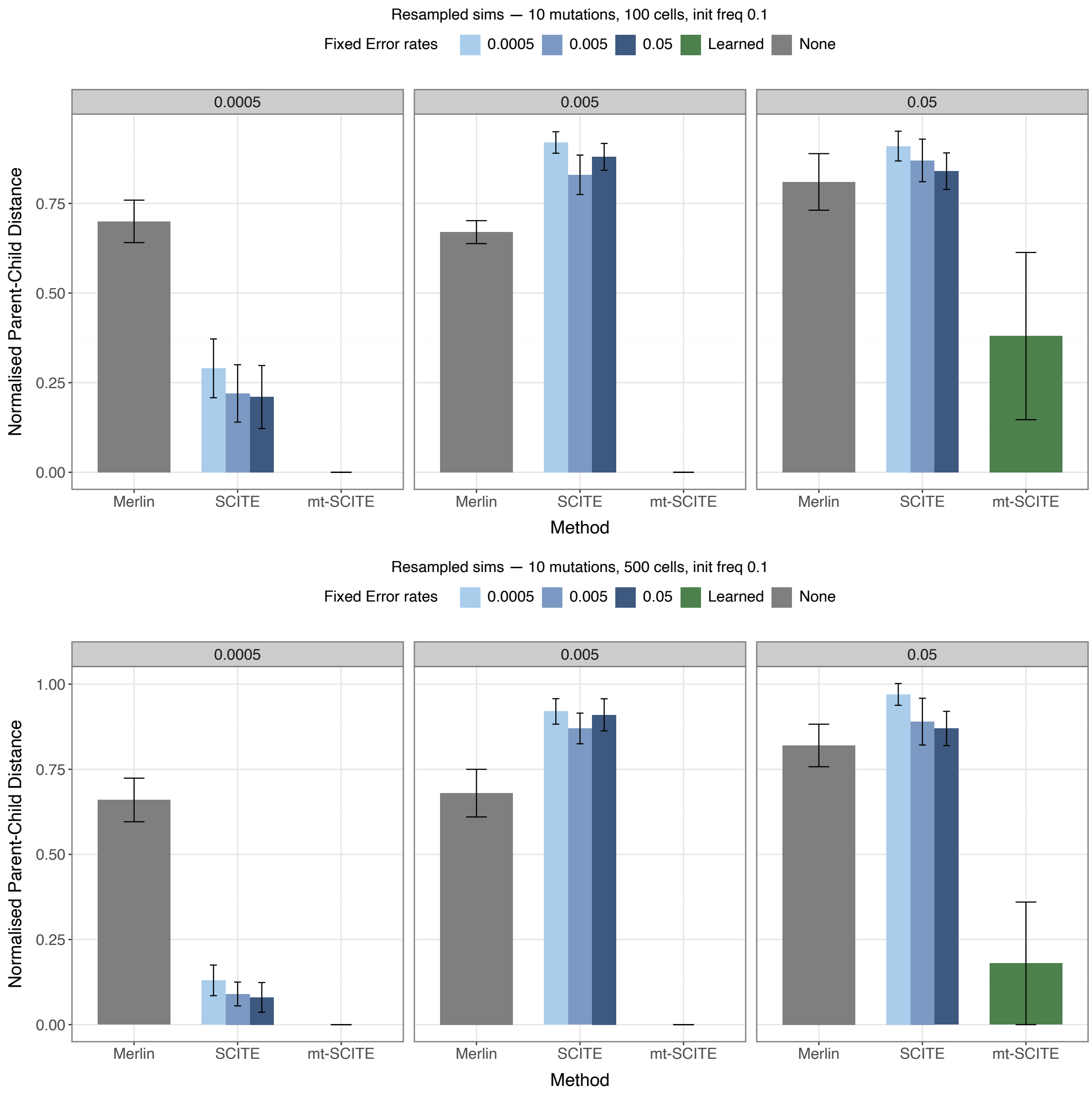


**Fig. 18**: Parent child distances for simulations with cell heteroplasmy level resampling are shown for 10 mutations, an initial mutation frequency of 10%, and (a) 100 cells and (b) 500 cells.

#### Small coverage

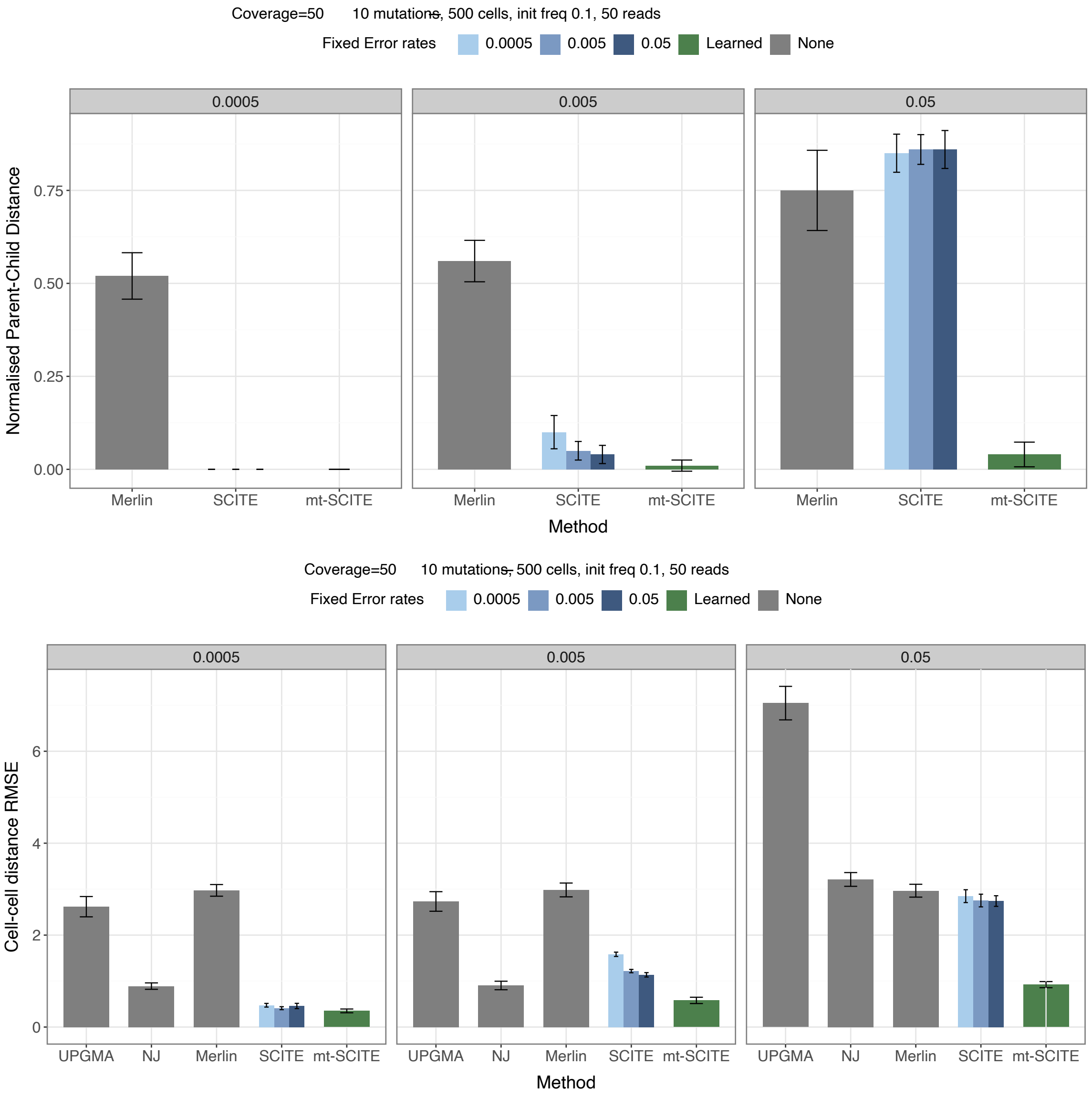


**Fig. 19**: Performance for coverage of 50 reads in terms of (a) parent child and (b) cell-cell distance for 10 mutations, an initial mutation frequency of 10%, and 500 cells.

For (b) we reuse the existing simulation framework with sequencing coverage reduced to 50 reads, which is more realistic for scRNA-seq data. mt-SCITE retains strong performance and remains the most accurate method across error rates (Fig. 19), consistent with coverage being an explicit part of its observation model, so that the support for a mutation is weighed appropriately at low depth.

#### Recurrent mutations

For (c), we test robustness to a violation of the infinite-sites assumption (ISA) by introducing a recurrent mutation. We first simulate a mutation tree for $n$ mutations as described above (Methods). Then we introduce an additional node representing the recurrent mutation as follows. We select a mutation node to be duplicated, $d$, from the tree at random. We create its duplicate $d'$ and search for a parallel lineage in the tree where to attach it to as a recurrent mutation. Thus we search for an attachment node $a$ uniformly at random all the nodes that are not $d$, ancestors of $d$ or a descendants of $d$; equivalently, $a$ lies in a parallel clade to $d$. Node $d'$ inherits its heteroplasmy frequency from its parent $a$ using the same Beta sampling procedure as described before, except the duplicated mutation that is shares with node $d$ is assigned a fixed heteroplasmy frequency. Cells then attach uniformly at random to the augmented tree and read counts are generated as in the main simulation.

We report results for 500 cells, an initial mutation frequency of 10%, and recurrent-mutation frequencies of 1% and 7% (Supplementary Figures 20, 21). mt-SCITE remains accurate on cell-cell distance recovery across all conditions. Mutation-tree reconstruction (parent-child distance) is likewise robust, degrading only when recurrence is frequent (7%) and the error rate is simultaneously high (0.05); at the lower recurrence rate (1%) performance is essentially unchanged from the infinite-sites case. mt-SCITE thus tolerates mild violations of the infinite-sites assumption, with accuracy affected only under the combination of recurrence at high VAF and high sequencing error.

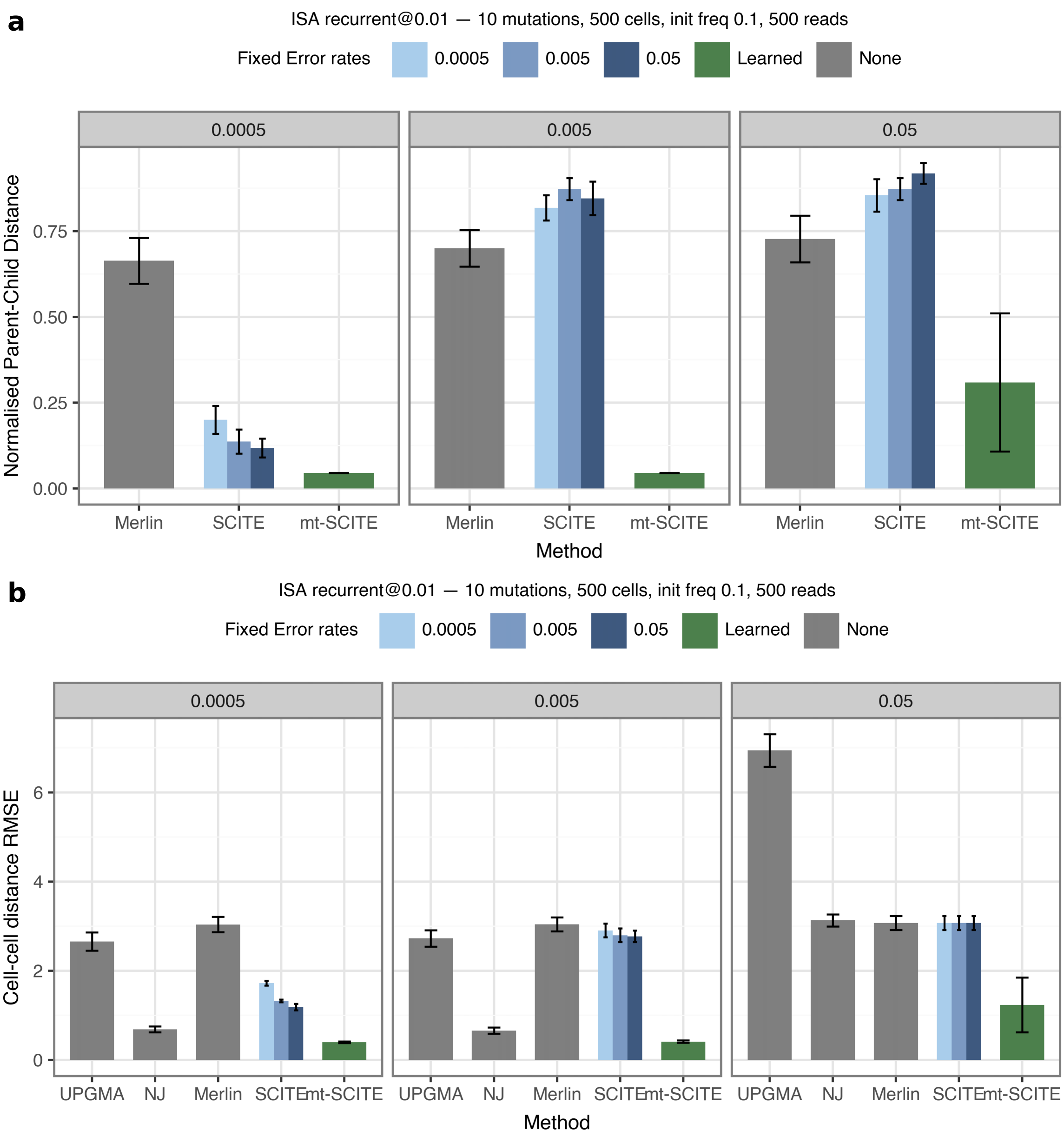


**Fig. 20**: Performance for recurrent mutation with VAF 1% for parent child (top) and cell-cell distance (bottom) for 10 mutations, an initial mutation frequency of 10%, and 500 cells.

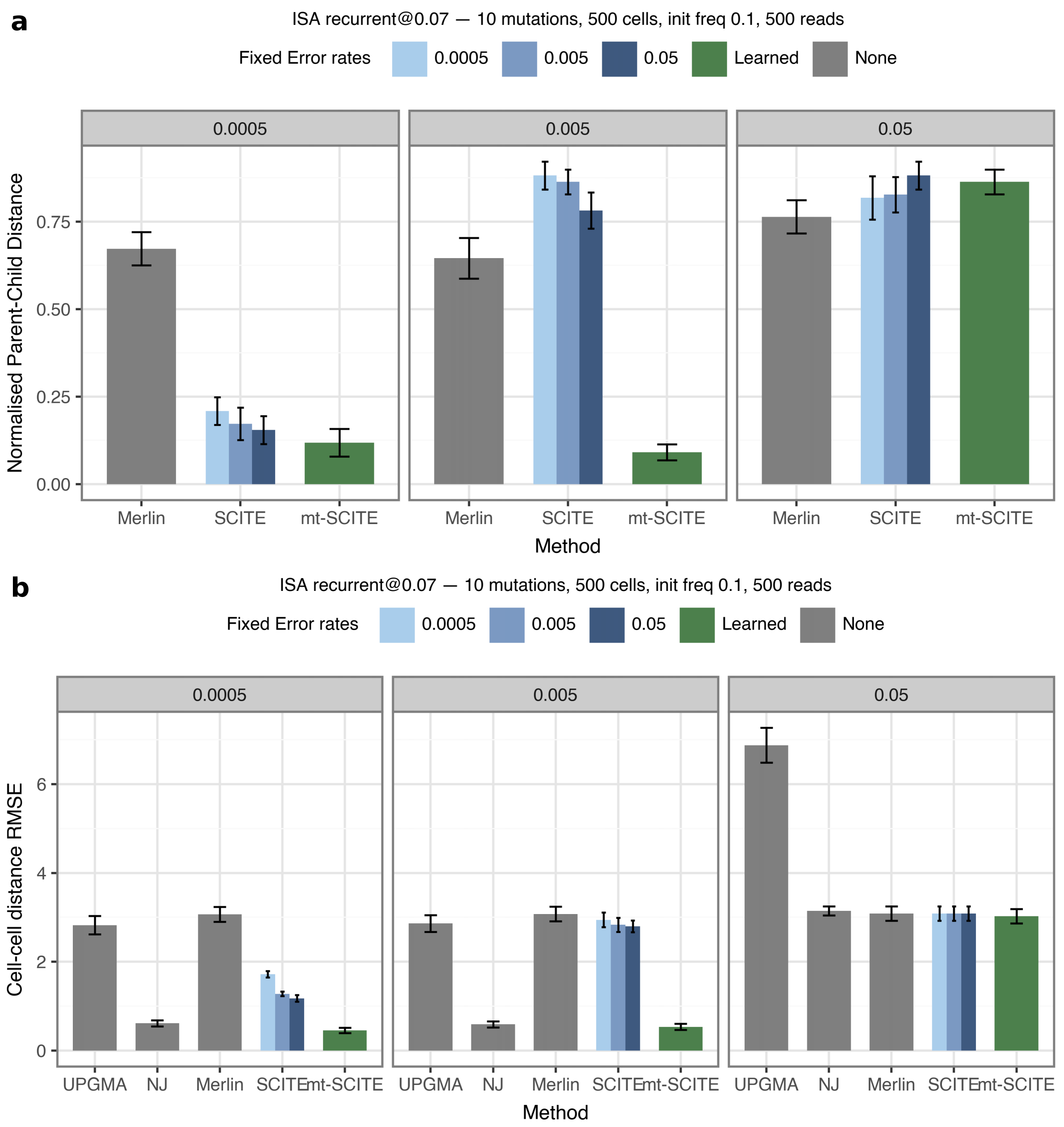


**Fig. 21**: Performance for recurrent mutation with VAF 7% for parent child (top) and cell-cell distance (bottom) for 10 mutations, an initial mutation frequency of 10%, and 500 cells.

#### Variant read count approximation

We approximate the term $P(v_{ij}^X \mid M_{ij} = X;\, c_{ij})$, the probability of observing $v_{ij}^X$ variant reads given that $X$ is the true state at coverage $c_{ij}$, with a uniform distribution over $1/c_{ij}$. In the main text, we draw this approximation directly, arguing for the uniform distribution as an uninformative prior.

Here, we motivate this approximation in a different and related way. Since the observed variant read count is a sample from the unknown heteroplasmy level $f \in [0, 1]$, we treat $f$ as latent and marginalise over it. We model the variant read count as $v_{ij}^X \mid f \sim \text{Binomial}(c_{ij}, f)$ with a non-informative prior $f \sim \text{Beta}(1, 1) = \text{Unif}(0, 1)$:

$$P\big(v_{ij}^X \mid M_{ij} = X;\, c_{ij}\big) = \int_0^1 \binom{c_{ij}}{v_{ij}^X} f^{\,v_{ij}^X} (1-f)^{\,c_{ij} - v_{ij}^X}\, \mathrm{d}f \tag{1}$$

$$= \frac{1}{c_{ij}+1} \int_0^1 \frac{f^{\,v_{ij}^X} (1-f)^{\,c_{ij} - v_{ij}^X}}{B(v_{ij}^X + 1,\, c_{ij} - v_{ij}^X + 1)}\, \mathrm{d}f = \frac{1}{c_{ij}+1} \tag{2}$$

$$\approx \frac{1}{c_{ij}}, \tag{3}$$

where the factor $1/(c_{ij} + 1)$ follows from $\binom{c_{ij}}{v_{ij}^X} = (c_{ij} + 1)^{-1}\, B(v_{ij}^X + 1,\, c_{ij} - v_{ij}^X + 1)^{-1}$ and the remaining integrand is a normalised Beta density. The exact result is uniform over the $c_{ij}+1$ possible counts $v_{ij}^X \in \{0, \ldots, c_{ij}\}$, which for typical coverage we approximate by $1/c_{ij}$.

### Beta distribution

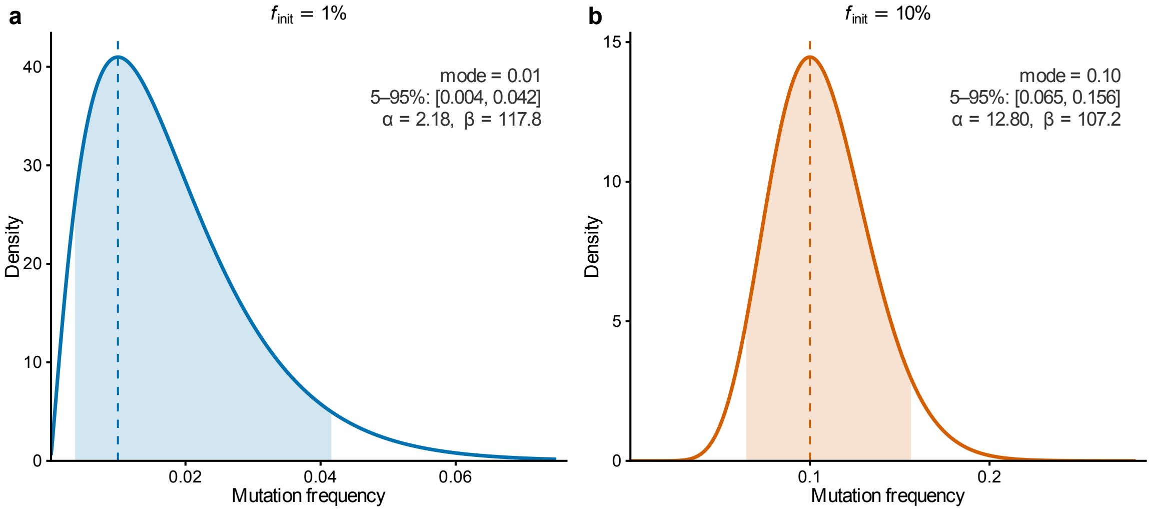


**Fig. 22**: Beta distribution used to sample mutation frequencies. Density of the Beta($f_{\text{init}}$, $\kappa = 120$) distribution for the two initial mutation frequencies used in the simulations, (a) 1% and (b) 10%. Dashed line shows the mode and the shaded region the 5–95% interval. $\alpha$ and $\beta$ are the corresponding Beta distribution shape parameters.

#### Raw variant allele frequencies at learned error rates

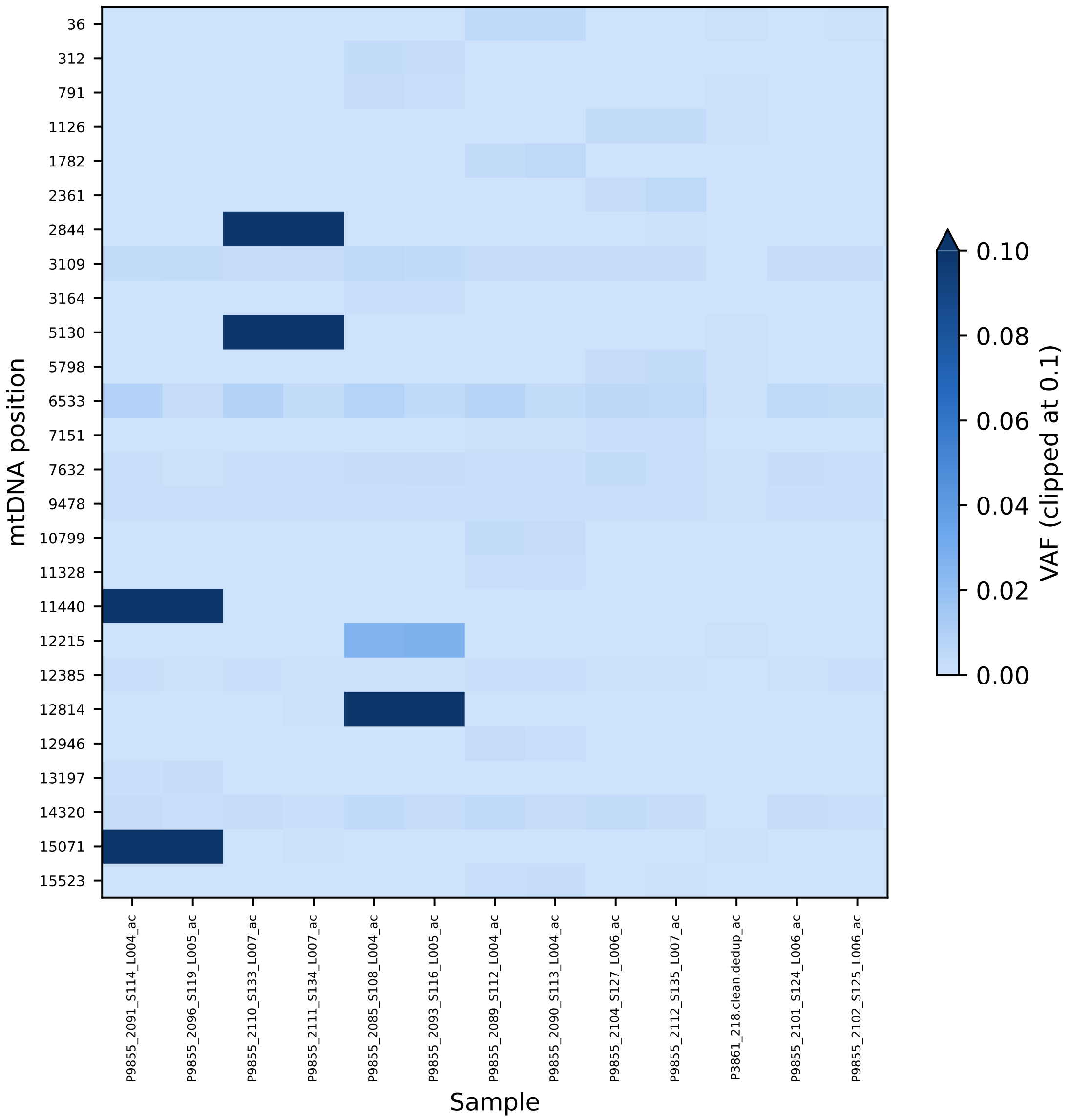


**Fig. 23**: Raw variant allele frequencies for mtDNA sites retained by mt-SCITE using its inferred error rate (0.0008), shown for the bulk ATAC sequenced T cell expansion dataset (Fig. 4).

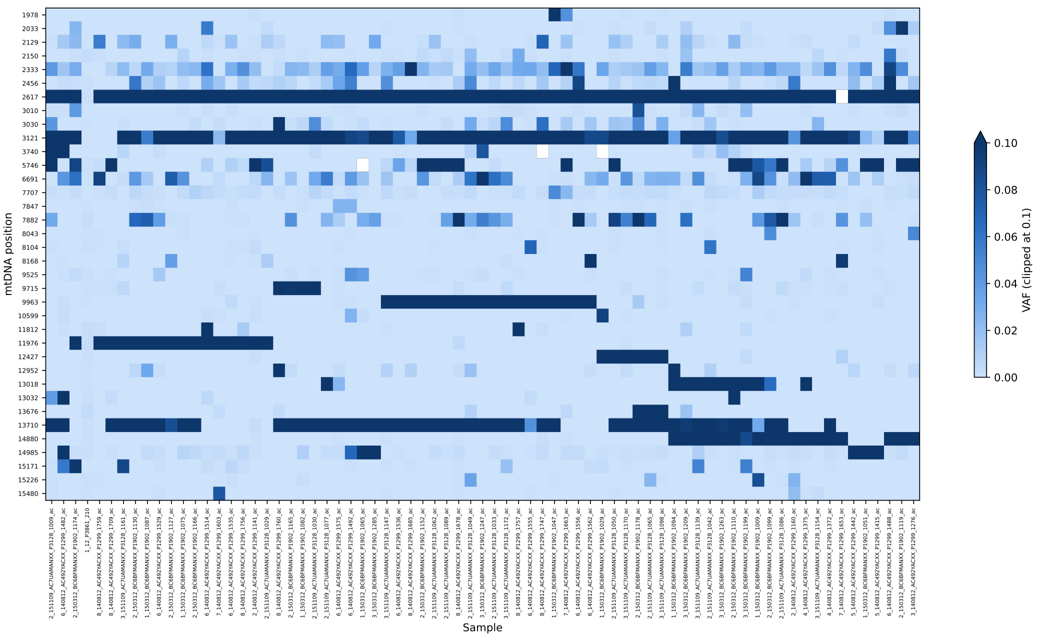


**Fig. 24**: Raw variant allele frequencies for mtDNA sites retained by mt-SCITE using its inferred error rate (0.06), shown for the T cell expansion dataset profiled by single-cell RNA-seq (Fig. 5).
